# Copper transport shapes parasite metabolism and dissemination in *Toxoplasma gondii*

**DOI:** 10.64898/2026.04.29.721613

**Authors:** Lucas Pagura, Jaden Todd-Nelson, Capucine Merlet, Jack C. Hanna, Laura Faulds, Sambamurthy Chandrasekaran, Shinhye Chloe Park, Kiana Koehler, Nishita Karikatti, Yamil Sanchez-Rosario, Michael D.L. Johnson, Alexandre Bougdour, Mohamed-Ali Hakimi, Caroline Bissardon, Peter Cloetens, Matthew W. Bowler, Stephen J. Fairweather, Giel van Dooren, Anita A. Koshy, Clare R. Harding

## Abstract

Copper is a conserved cofactor; required for aerobic mitochondrial respiration by eukaryotes. In parasitic eukaryotes the mechanisms of copper uptake, and the role of copper within the cell, are not well understood. Here, we identify and validate two copper transporters of *Toxoplasma gondii* (Ctr1 and Ctr2) and determine the role of copper in parasite metabolism, dissemination and pathogenesis. We show Ctr1 is localised to the plasma membrane where it is required for copper uptake. Deletion leads to a loss of parasite-associated copper, linked with a significant growth defect and dramatic metabolic shift from mitochondrial respiration to glycolysis. Loss of Ctr1 was fully and rapidly rescued by exogenous copper, demonstrating a potential secondary copper uptake pathway. In contrast, Ctr2 localised to intercellular vesicles and was dispensable during rapid tachyzoite growth. Ctr2 expression was important for parasite survival under copper-limited conditions and for parasite survival and dissemination during *in vivo* infection. Together, these findings reveal a critical role for copper uptake in shaping metabolic plasticity and highlight the importance of nutrient transport in apicomplexan metabolism and pathogenesis.

## Introduction

The fundamental chemical properties of transition metals, including copper, iron, and zinc, power cellular energetics. These same properties make transition metals potentially harmful, leading to the evolution of highly specialised and tightly regulated processes for metal acquisition, storage and usage within cells. Of these metals, copper is the least abundant in cellular contexts. This is due to its high redox activity and stability in the Irving-Williams series, meaning that excess copper leads to mismetallation of essential enzymes and cell death^1,2^. These redox properties are exploited by cells to drive electron transfer in a range of key reactions across the domains of life, including respiration, redox control, pigmentation and neuropeptide production^4^. This functional duality presents a challenge to intracellular pathogens which must acquire and utilise copper from their intracellular environment, despite host-driven restriction and redistribution of copper, both locally and systemically^5,6^.

*Toxoplasma gondii* is a highly successful intracellular parasite, capable of infecting all warm-blooded animals, including up to 30% of the human population. Although typically asymptomatic, infection can have devastating pathological consequences in immunocompromised individuals and in the developing foetus^7,8^. During acute infection, *Toxoplasma* tachyzoites subvert host cell nutrients (including sugars, lipids, amino acids and transition metals) to power rapid division and dissemination^9–12^. The availability of parasite nutrients differs across the cell types and tissues *Toxoplasma* infects, resulting in a range of potential nutritional environments, and potentially promoting the metabolic flexibility of these parasites^13,14^. Faced with diverse environmental stresses (e.g. pH, nutrient starvation, host immune pressure), a small number of parasites differentiate into slow-dividing bradyzoites, which develop within intracellular tissue cysts. Within these tissue cysts, parasites are largely metabolically quiescent and appear resistant to both host immune responses and currently available medical therapies^15^.

In the related apicomplexan parasite *Plasmodium,* copper is required in the blood and insect stages^16,17^, although copper’s roles were not defined. The most established function of copper is as a cofactor for cytochrome c oxidase, also known as complex IV in the mitochondrial electron transport chain (ETC). Complex IV contains two copper centres (CuA and CuB) which serve to shuttle electrons, generated from mitochondrial dehydrogenases and complexes II and III, to the terminal acceptor (oxygen) generating a proton gradient that drives ATP production via ATP synthase (also called complex V)^18^. Although there are significant lineage-specific features in the ETC in the Apicomplexa^19–21^, recent structures confirm the conservation of copper-binding in *Toxoplasma* complex IV^19^ and conservation of the mitochondrial copper chaperones^22,23^. Previous studies have shown the apicomplexan mitochondrial ETC is required for mitochondrial ATP production and pyrimidine biosynthesis and its activity is important in maintaining *Toxoplasma* fitness^19,24–26^. Beyond complex IV, a predicted copper amine oxidase of unknown function is encoded, which genome-wide knockout screens suggest is not required for parasite fitness^27–29^. BLAST searches suggest that *Toxoplasma* encodes no other predicted cuproenzymes.

Cellular copper import is mediated by the highly conserved SLC31 (Ctr) family of copper transporters. Ctr proteins trimerize in cellular membranes, forming a channel that transports Cu^+^ with high selectivity^30,31^. In many organisms, including mammalian cells and yeast, Ctr1 functions as a high-affinity plasma membrane copper importer, whereas Ctr2 has lower affinity and localises to the endolysosome system, where its roles are still debated^32,33^. Within the Apicomplexa, the only previous study of copper transporters demonstrated that PF3D7_1421900, (putative PfCtr1) localises to the parasite periphery and binds copper via its N-terminus^34^. However, copper transporters and the role of copper have not been assessed in *Toxoplasma gondii*.

Here, we identify and functionally characterize the two major copper transporters in *Toxoplasma gondii,* Ctr1 and Ctr2. We show Ctr1 localises to the parasite plasma membrane and validate its transport activity in exogenous expression in *Xenopus* oocytes. Ctr1-mediated copper import is essential for mitochondrial respiration via complex IV activity, and that its loss drives a substantial metabolic rewiring away from mitochondrial respiration in the parasite. Loss of Ctr1 can be fully and rapidly rescued by addition of exogenous copper. In contrast, Ctr2 is dispensable in tachyzoites in copper-replete environments but is copper-responsive and required in copper-depleted environments *in vitro*. We further show that Ctr2 is required for dissemination during acute infection *in vivo,* consistent with the *in vivo* environment restricting access to this essential nutrient.

## Results

### Toxoplasma gondii encodes putative copper transporters

Copper uptake in eukaryotes is mediated by the highly conserved SLC31 (Ctr) family of proteins. We examined the *Toxoplasma* genome and identified three genes predicted to encode Ctr domains TGME49_262710 (Ctr1), TGME49_249200 (Ctr2), and TGME49_221350 (Ctr3). Phylogenetic analysis of Ctr1 and Ctr2 demonstrated that both are conserved among Coccidia (**Fig. 1a**) and cluster together with the predicted *Plasmodium ctr1* and *ctr2*, with *ctr2* having undergone less divergence between the Coccidia and Haemosporida. All three copper transporters encode at least three transmembrane domains and the conserved and essential MxxxM (Mx_3_M) and GxxxG (Gx_3_G) motifs in the second and third transmembrane domains, respectively **(Fig. 1b)**. The Mx_3_M motif coordinates copper binding and trimerization. Each methionine interacts across subunits, generating a double barrier that is critical for Cu^+^ selectivity, critical for copper transport^31,34–36^. The Gx_3_G domain has been reported to be essential for the trimeric assembly interface between subunits^30,37^. In addition, *Toxoplasma* Ctr1 and Ctr2 also share a highly conserved sequence of unknown function^36^ (MPMxFX_8_LF) in the N-terminal extension, predicted to sit on top of the channel formed in the trimeric structure, however this is absent from Ctr3. Both *ctr1* and *ctr2* are highly expressed in tachyzoites and bradyzoites, with maintenance of *ctr1* expression throughout the parasite lifecycle. In contrast, *ctr3* shows minimal expression at any lifecycle stage (**Fig. 1c**).

**Figure 1.**
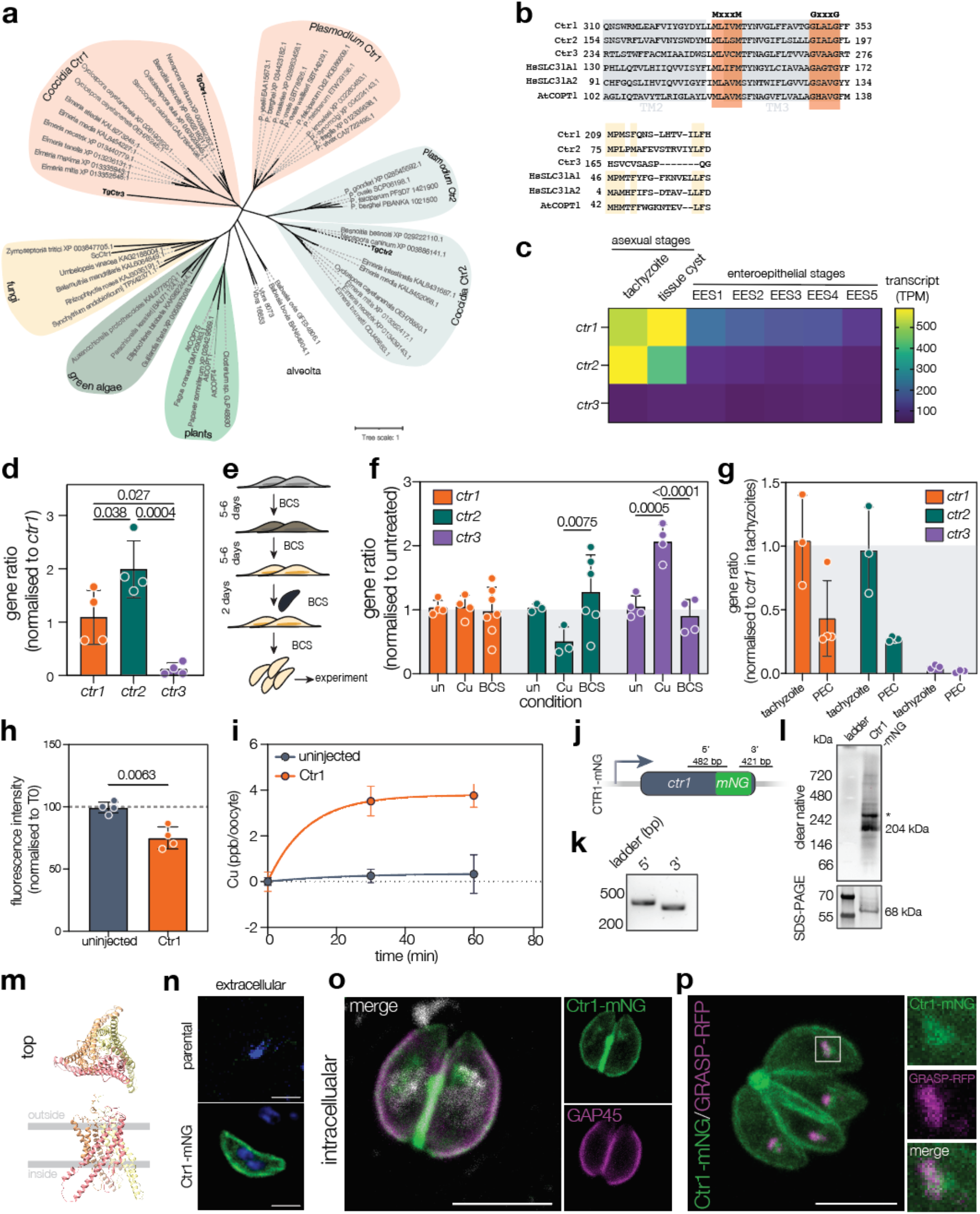
*Toxoplasma gondii* encodes highly conserved copper transporters from the Ctr family. **a.** Cladogram showing how Ctr1, 2 and 3 are evolutionarily related to other coccidian Ctr proteins. **b.** Alignment of Ctr1 (TGME49_262710), Ctr2 (TGME49_249200) and Ctr3 (TGME49_221350) along with human homolog proteins HsSLC31A and HsSLC31A2 and *Arabidopsis thaliana* AtCOPT1. Ctr protein showing conserved MxxxM and GxxxG (orange) motifs, transmembrane regions 2 and 3 highlighted in grey. **c.** Heat map showing *ctr* transcript levels during the different stages of *T. gondii* life cycle. **d.** Quantification of *ctr1*, *ctr2* and *ctr3* transcripts, normalized to actin and *ctr1* by qRT-PCR. Bars are at mean ± SD of three independent experiments. *p* values from one-way ANOVA with Tukey’s correction. **e.** Schematic of BCS (100 µM) treatment for copper depletion. **f.** Quantification of *ctr1*, *ctr2* and *ctr3* transcripts, normalized to actin as housekeeping control, in untreated, copper treated or Cu-depleted parasites. Bars are at mean ± SD of three independent experiments. *p* values from two-way ANOVA with Tukey’s correction. **g.** Quantification of *ctr1*, *ctr2* and *ctr3* transcripts, normalized to actin as housekeeping control, in tachyzoites and peritoneal exudate 5 days post infection (dpi) (PEC), normalized to *ctr1* in tachyzoites. Bars are at mean ± SD of three independent experiments, *p* values from one-way ANOVA with Tukey’s correction. **h.** Fluorescence intensity of PhenGreen dye from control or Ctr1-expressing *Xenopus laevis* oocytes after incubation with 20 µM Cu^+^ for 60 min. Bars are at mean ± SD of four independent experiments. *p* values from unpaired two-tailed *t* test with Welch’s correction. **i.** Copper uptake over time by control or Ctr1-expressing *Xenopus laevis* oocytes after 30 or 60 min of incubation with 25 µM Cu^+^ measured by ICP-MS. Bars are at mean ± SD of 5 independent samples. **j.** Schematic for endogenous tagging of Ctr1 with mNeonGreen. **k.** Integration PCR confirming correct integration of mNeonGreen. **l.** Western blot from a clear-native PAGE (top) where a signal corresponding to the trimer is observed (204 kDa), and from a SDS-PAGE (bottom), where signal for the monomer is observed at the expected size (68 kDa). **m.** AlphaFold modeling of Ctr1 trimer. **n.** Ctr1-mNG is seen at the periphery of the cell in extracellular parasites, no signal from the parental line. Scale bar 5 µm. **o.** Ctr1-mNG co-localises with the peripheral marker GAP45, and extends over the apical and basal caps, indicating plasma membrane localisation, DAPI is in white. Scale bar 5 µm. **p**. Ctr1-mNG parasites transiently expressing GRASP-RFP show overlapping signal with internal Ctr1-mNG signal. Scale bar 5 µm.

### *Toxoplasma* copper transporters are transcriptionally regulated by copper

We first assessed the expression of the proposed copper transporters in RH tachyzoites using qRT-PCR. We found that *ctr2* shows significantly higher expression than *ctr1* (one-way ANOVA, Tukey’s correction) while, in support of the RNAseq data above, ctr3 was barely above the limit of detection (**Fig. 1d**). To determine if the proposed copper transporters were transcriptionally responsive to changing copper levels, we performed qRT-PCR after treatment. Parasites were either treated with 5 µM CuSO_4_ for 24 h, or host cells were grown in the presence of the copper chelator Bathocuproinedisulfonic acid (BCS)^2^ at 100 µM for two passages (10-14 days), then *Toxoplasma* was grown for two passages (36-42 h) in BCS-treated host cells (**Fig. 1e**). *ctr1* (normalised to parasite actin as a housekeeping control) showed no change in transcription under the copper conditions tested, however *ctr2* was significantly downregulated upon copper treatment (*p* = 0.0075, two-way ANOVA, Tukey’s correction) while *ctr3* was only significantly (*p* < 0.0001, two-way ANOVA, Tukey’s correction) upregulated (**Fig. 1f**). In contrast, the iron transporter *vit*^38^ did not show any significant changes under any of the conditions tested (**Fig. S1a**). We also determined the expression of all three genes in vivo. Mice were infected with the cystogenic Pru line to limit lethality, and peritoneal exudate (PEC) cells were collected at 5 days post infection (dpi). qRT-PCR was performed and normalised to tachyzoites grown *in vitro*. Here, we see both *ctr1* and *ctr2* were downregulated in infection. *ctr3* was barely expressed under both tachyzoite and *in vivo* conditions (**Fig. 1g**).

### Ctr1 is a functional copper transporter

To confirm the activity of Ctr1 and Ctr2, we attempted to ectopically express both in *Xenopus laevis* oocytes, as previously described^9,39^. Surface expression of Ctr1-HA was confirmed by streptavidin pulldown of biotinylated surface proteins, followed by western blotting (**Fig. S1b**). Although Ctr2-HA was detected by WB in whole protein extracts from oocytes (**Fig. S1**c), surface biotinylation assays did not retrieve the protein (**Fig. S1b**), suggesting that Ctr2-HA is retained in intracellular compartments, making further characterisation by this assay unfeasible. To determine if expression of Ctr1 was sufficient to transport Cu in oocytes, Ctr1-expressing and control oocytes were incubated in the presence of 20 µM reduced Cu^+^ for 60 min. All oocytes were then injected with the dye PhenGreen, whose fluorescence is quenched in the presence of cations, including copper^40^, and the fluorescent signal quantified. No change (compared to time 0) in fluorescence could be seen in control oocytes, demonstrating no loss of membrane integrity over the course of the experiment. However, incubation of Ctr1-HA-expressing oocytes with Cu^+^ for 1h led to a significant (*p* = 0.0063, unpaired two-tailed *t* test with Welch’s correction) decrease in PhenGreen signal (**Fig. 1h**), suggesting cation uptake under these conditions.

To confirm Ctr1 activity, we incubated Ctr1-HA expressing or control oocytes with 25 µM Cu^+^ for 0, 30 and 60 min and quantified copper uptake using ICP-MS. We saw an increase in copper content of Ctr1-expressing oocytes, not observed in the control oocytes over time (**Fig. 1i**), confirming the ability of Ctr1 to transport Cu^+^ across the membrane.

These data confirm that *Toxoplasma* expresses copper-responsive putative copper transporters, and that plasma membrane-localised Ctr1 can transport copper upon ectopic expression.

### Ctr1 is localised to the parasite plasma membrane

As initial attempts to endogenously C-terminally tag Ctr1 were unsuccessful, we decided to add a mNeonGreen (mNG) fluorescent tag 2aa before the stop codon, allowing preservation of the 3’ UTR codon (**Fig. 1j**). We confirmed correct integration of the tag by PCR (**Fig. 1k**). Western blotting under denaturing conditions allowed detection of a band of the expected size (68 kDa). Under native condition we were instead able to detect a signal at the expected size of a trimer (204 kDa) as well as a higher band of unknown composition (**Fig. 1l**). This is consistent with previous reports that Ctr1 proteins functioning as trimers^30,37^ and is further supported by AlphaFold models of Ctr1 (**Fig. 1m**) and Ctr2 and Ctr3 (**Fig. S1d**). We were able to localise the majority of Ctr1-mNG to the plasma membrane in both extracellular (**Fig. 1n**) and intracellular parasites (**Fig. 1o**). We also observed accumulation of signal within the parasites, which partially co-localised with the cis-Golgi marker GRASP-RFP^41^ (**Fig. 1p**) but not with the plant-like vacuolar compartment (PLVAC) marker CPL (**Fig. S1e**), suggesting that Ctr1 is trafficked to the plasma membrane via the Golgi. Unlike the iron and zinc transporter ZFT^9^, we did not see changes in localisation dependent on the number of parasites/vacuole (**Fig. S1f**). To determine if the localisation of Ctr1 was dependent on the copper conditions, we imaged Ctr1-mNG after treatment with high copper (20 or 50 µM CuSO_4_) or low copper (BCS-treated) for 24 h. However, we did not observe any significant changes in localisation or abundance over these conditions (**Fig. S1g**).

### Deletion of *ctr1* impairs tachyzoites growth

Based on the data above, we investigated the role of Ctr1 in *Toxoplasma*. We used the CRISPR/Cas9 system to generate a transgenic knockout of *ctr1* in the RHΔKu80:tdTomato red-fluorescent parasite line^38^ by replacing the native *ctr1* sequence with the pyrimethamine resistance gene *dhfr* (**Fig. 2a**). Surprisingly, although *ctr1* was predicted to be fitness conferring from a genome-wide screen (phenotype score -2.62^29^), we successfully generated a full knockout of this gene, as confirmed by integration PCR (**Fig. S1h and i**). We confirmed loss of *ctr1* by qRT-PCR and found no significant change in expression of *ctr2* or *ctr3* upon knockout of *ctr1* (**Fig. S1j**).

**Figure 2.**
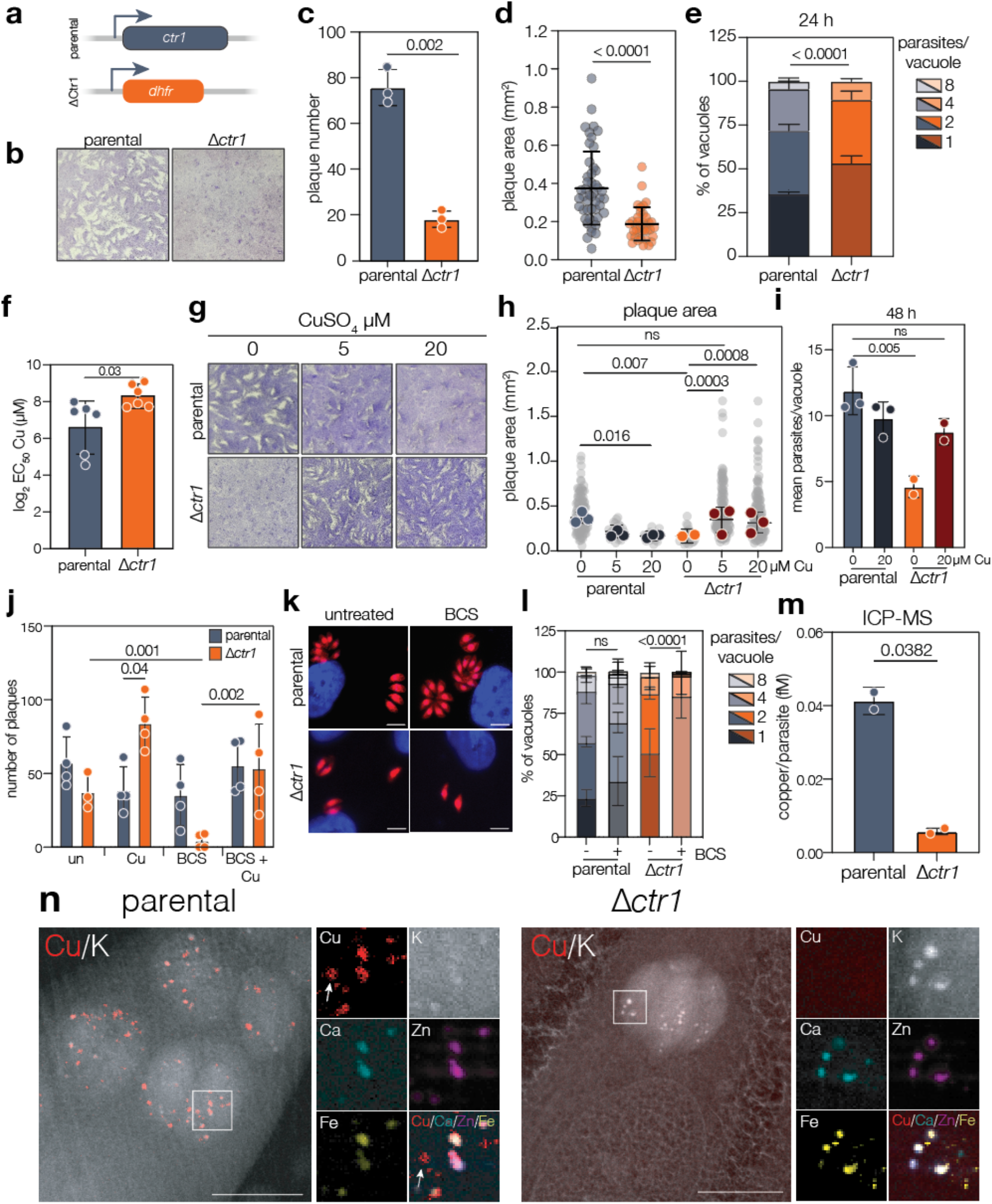
Loss of *ctr1* impairs parasite growth, a phenotype that is rescued by exogenous copper. **a.** Schematic of Δ*ctr1* strain construction by replacing the native *ctr1* sequence by pyrimethamine resistance gene *dhfr*. **b.** Plaque assay of parental and Δ*ctr1* parasites in regular conditions. Quantification of plaque number (**c**) and area (**d**) for parental and Δ*ctr1* parasites. Bars are at mean ± SD*. p* values are from unpaired two-tailed *t* test with Welch’s correction. **e.** Quantification of percentage of parasites/vacuole in parental and Δ*ctr1* parasites after 24 h of infection. 100 vacuoles for each condition were counted. Bars at mean ± SD of three independent replicates. **f.** Copper EC_50_ determined after 4 days of incubation, showing reduced susceptibility for Δ*ctr1* parasites compared to the parental line. Bars are at mean ± SD*. p* value from unpaired two-tailed *t* test with Welch’s correction. **g.** Plaque assay of parental and Δ*ctr1* parasites untreated and CuSO_4_ treated. **h.** Quantification of plaque area generated by parental or Δ*ctr1* parasites under different copper treatment. *p* values are from two-way ANOVA with Sidak correction. **i.** Mean parasites/vacuole in parental and Δ*ctr1* parasites after 48 h of infection in untreated or 20 µM CuSO_4_ supplemented condition. Bars are at mean ± SD of three independent replicates. *p* values from one-way ANOVA with Tukey correction. **j.** Quantification of plaque number of parental and Δ*Ctr1* parasites after treatment with 5 µM CuSO_4_, 100 µM BCS or combination of both. Bars are at mean ± SD. *p* values from two-way ANOVA with Tukey correction. **k.** Fluorescence images of parental and Δ*ctr1* parasites untreated or treated with 100 µM BCS for 48 h. Scale bars 5 µm. **l.** Quantification of percentage of parasites/vacuole in parental and Δ*ctr1* parasites after 48 h in regular or BCS copper depleted condition. 100 vacuoles for each condition were counted. Bars at mean ± SD of three independent replicates. *p* value from two-way ANOVA with Tukey correction. **m.** ICP-MS quantification of Cu/parasite (fM) from parental and Δ*Ctr1* parasites. Bars are at the mean ± SD of three independent experiments. *p* values from unpaired two-tailed *t* test with Welch’s correction. **n.** XFM elemental maps of parental and Δ*ctr1* parasites showing copper (red) and potassium (white). Insets show overlap of copper, calcium, zinc, iron and potassium in foci in the parental line, although copper is also present with no overlap (arrows) in the parental line. Scale bar 5 µm.

Supporting the negative phenotype score, Δ*ctr1* parasites showed highly impaired growth by plaque assay (**Fig. 2b**), with significantly fewer (*p* = 0.0021, unpaired two-tailed *t* test with Welch’s correction) (**Fig. 2c**) and smaller (*p* < 0.0001, unpaired two-tailed *t* test with Welch’s correction) (**Fig. 2d**) plaques after 7-8 days. To determine if this defect is due to a defect in intracellular replication, we quantified parasites/vacuole at 24 h post infection and found a significant increase in the number of vacuoles containing one parasite and a decrease in the number of vacuoles containing four parasites (*p* < 0.0001, *p* = 0.0004 respectively, two-way ANOVA with Sidak correction), demonstrating that in the absence of *ctr1*, parasites are deficient in replication (**Fig. 2e**).

We predicted that lack of copper uptake could protect against excess copper toxicity. To test this, parental and Δ*ctr1* parasites were treated with increasing concentrations of copper and total parasite fluorescence (as a proxy for parasite replication^38^) was measured after 4 days. Absence of *ctr1* significantly (*p* = 0.03, unpaired two-tailed *t* test with Welch’s correction) protected against copper toxicity (ΔEC_50_ = 193 µM) (**Fig. 2f**), supporting the hypothesis that *ctr1* mediates parasite copper internalisation.

In other systems, loss of a metal transporter can be complemented by addition of exogenous substrate^42,43^. To determine the effect of copper supplementation in the absence of *ctr1*, we performed plaque assays in the presence of exogenous copper. The addition of low copper concentrations (5-20 µM) appears toxic to the parental line, significantly (*p* < 0.02, two-way ANOVA with Sidak correction) reducing both the number and size of plaques formed (**Fig. 2g and h, Fig. S1k**). However, addition of copper enhanced growth of Δ*ctr1* parasites, allowing the parasites to form more (**Fig. S1k**) and significantly *(p* = 0.0008, two-way ANOVA with Sidak correction) larger plaques, in a concentration-dependent manner (**Fig. 2g and h**). In contrast, no rescue in parasite growth (as measured by plaque area) was observed upon supplementation with iron (ferric ammonium chloride) or zinc (zinc sulphate) (**Fig. S1l**), confirming the specificity of this effect. To validate the copper-mediated rescue, we quantified mean parasite number/vacuole at 48 h post infection. As expected, deletion of *ctr1* led to a significant (*p* = 0.005, one-way ANOVA with Tukey correction) decrease in average parasite number/vacuole, but addition of 20 µM copper increased the average parasites/vacuole almost to the parental levels (**Fig. 2i**). These data demonstrate that the growth phenotype resulting from loss of *ctr1* could be rescued by exogenous copper.

While the growth defect caused by the absence of *ctr1* was severe, the parasites were viable and could be maintained in culture. To determine the effect of loss of *ctr1* in a low copper environment, we grew parental and Δ*ctr1* parasites in BCS-treated host cells for one passage and then performed a plaque assay in BCS-treated host cells. BCS treatment did not significantly affect plaque number of the parental line (**Fig. 2j**), however we saw almost no plaque formation in BCS-treated conditions in the Δ*ctr1* parasite line, although this was rescued (*p* = 0.002, two-way ANOVA, Tukey correction) upon addition of 5 µM copper (**Fig. 2j**), demonstrating the copper-specificity of BCS treatment. We also quantified the number of parasites/vacuole under these conditions and found that copper depletion by BCS-treatment exacerbated the growth defect in the Δ*ctr1* parasite line, with significantly more single parasite vacuoles (*p* < 0.0001, two-way ANOVA with Tukey correction) in BCS-treated cells than in untreated ones (**Fig. 2k and l**).

We hypothesized that if *ctr1* is the major parasite copper transporter, its deletion would impact parasite-associated copper levels. To test this, we quantified total parasite-associated metals using ICP-MS. Although we did not see any significant change in iron or zinc levels in the Δ*ctr1* line (**Fig. S1m** and **S1n**), we did see a significant (*p* = 0.038, unpaired two-tailed *t* test with Welch’s correction) decrease in parasite-associated copper in the Δ*ctr1* line (**Fig. 2m**), supporting our hypothesis that *ctr1* is required to maintain parasite copper levels.

To determine how deletion of *ctr1* impacted copper distribution within the parasites, we used high resolution X-ray fluorescence microscopy (XFM) on cryofixed intracellular parasites. In the parental line, we saw high concentrations of copper in multiple small foci, distributed throughout the parasite (**Fig. 2n**). Some of these copper foci overlapped with iron, zinc and calcium. Previous studies have localised calcium, iron and zinc vacuolar transporters to a lysosomal-like organelle called the PLVAC^38,44,45^, suggesting that copper is also stored within these dynamic structures. Interestingly, some copper foci did not overlap with any other detected elemental foci (arrows), however the identity and role of these remains unknown. XFM resolution can be limited by ice thickness during sample preparation, however, despite better sample preparation (as observed by the detailed host nuclear architecture in the potassium channel), in the Δ*ctr1* sample, we did not observe any detectable copper foci (**Fig. 2n**), although the localisation of other detected elements remained unchanged.

Together, these observations demonstrate that *ctr1* is the major copper transporter in tachyzoites of *T. gondii,* and that copper is important for parasite growth.

### Metabolomics reveals copper-dependent changes in pyrimidine biosynthesis and central carbon metabolism

To investigate why loss of *ctr1* led to this significant growth defect, we performed RNAseq on parental, Δ*ctr1* and Δ*ctr1* parasites supplemented with 600 µM copper (to ensure total complementation) for 24 h prior to collection. We found a transcriptional shift upon knockout of *ctr1*, with 596 genes downregulated (log_2_ fold change < -1.5, adj.p < 0.05) and 92 upregulated (log_2_ fold change > 1.5, adj.p < 0.05), compared to the parental line (**Fig. S2a** and **b**). This change in transcriptional profile was largely rescued by the addition of copper, which restored parental expression levels of 88% of the downregulated and 38% of the upregulated genes (**Fig. S2c**).

Despite these changes, examination of the transcriptomes did not reveal changes in specific pathways which could explain the significant growth defect. Given that the only known cuproenzyme in *Toxoplasma* functions in the electron transport chain, we hypothesised that metabolic changes, which were not reflected in the transcriptome, could be leading to the parasite’s growth defect. To examine this, we quantified metabolite abundance using untargeted, quantitative metabolomics. PCA analysis demonstrated good agreement between replicates and revealed a clear distinction between *Δctr1* parasites and the parental line (**Fig. S2d**), with significant (log_2_ fold change < -0.6 or > 0.6, adj.p < 0.05) accumulation of 89 metabolites and decreased abundance of 56 in the Δ*ctr1* parasites (**Fig. S2e**). Examination of altered metabolites revealed significant accumulation of pyrimidine intermediates (with accumulation of N-carbonyl aspartate, dihydroorotate and orotate) consistent with a block at the ubiquinone-dependent step catalysed by dihydroorotate dehydrogenase (DHODH). In contrast, we saw decreased levels of oxaloacetate and aspartate (**Fig. 3a** and **S2e**). While aspartate and oxaloacetate both act as precursors for pyrimidine biosynthesis^46^, oxaloacetate is a central TCA cycle intermediate, and aspartate is a proteinogenic amino acid with roles extending well beyond this pathway^46^. Unlike the related parasite *Plasmodium, Toxoplasma* can salvage pyrimidines from its environment which can rescue genetic ablation of pyrimidine biosynthesis^26,47^. To determine if disruption of pyrimidine biosynthesis was linked to the replication defect, we tested if exogenous uracil could rescue Δ*ctr1* growth. Addition of 250 µM uracil led to a small but significant (*p* < 0.0001, one-way ANOVA, Dunnett’s correction) increase in plaque size and in plaque number (*p* = 0.037, *t* test). Although the rescue of plaque number was comparable, the impact of uracil on plaque size (approx. 140% increase) was significantly less than the impact of copper (940%) (**Fig. S2f** and **S2g**).

**Figure 3.**
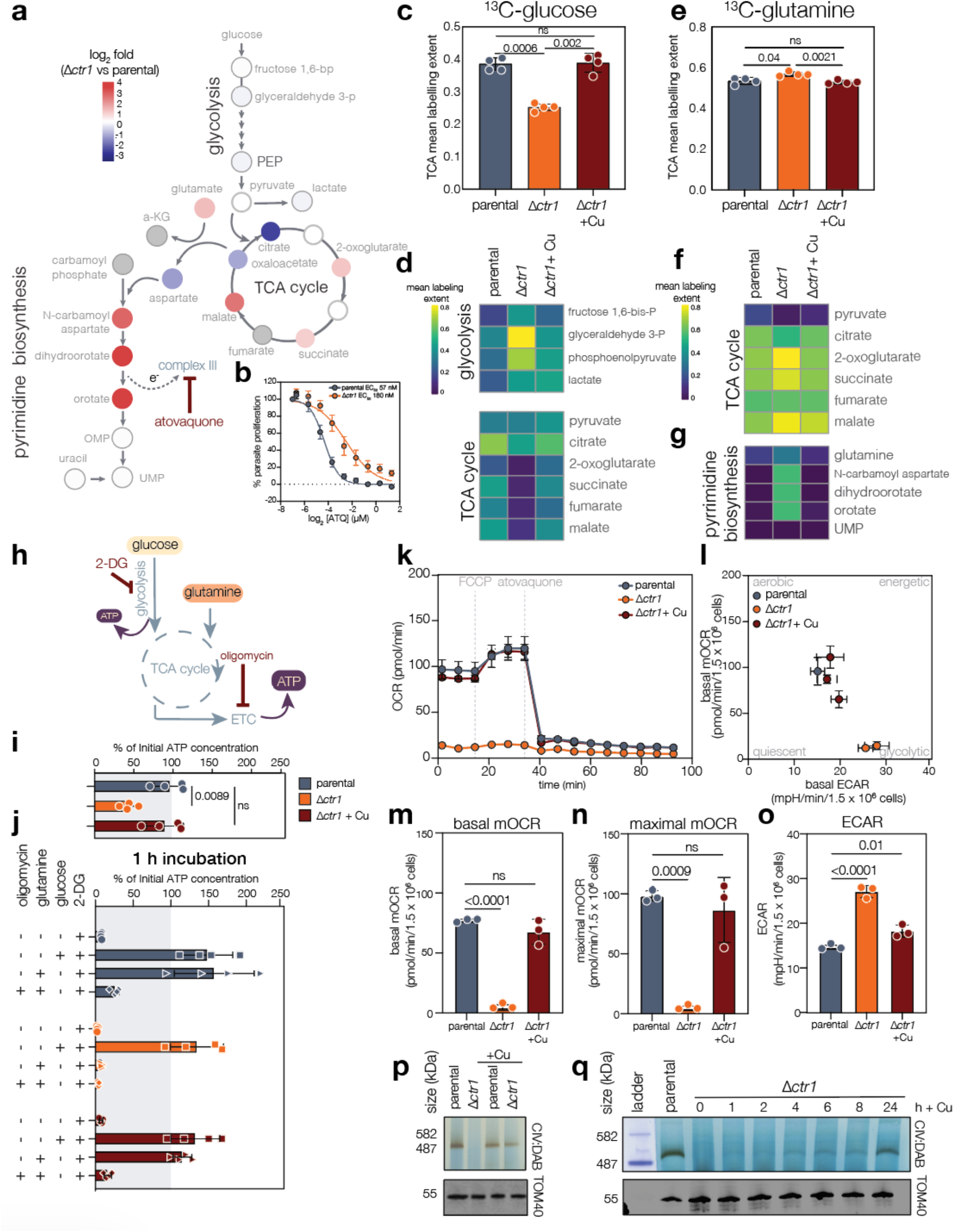
Deletion of *ctr1* significantly impacts parasite metabolism and mitochondrial respiration. **a.** Metabolic diagram of central carbon metabolism and pyrimidine biosynthesis. Point colour indicates fold change in abundance in Δ*ctr1* over parental parasites. **b**. Sensitivity of parental and Δ*ctr1* parasites to atovaquone, points mean of four independent experiments, ± SEM**. c.** Mean labelling extent of TCA associated metabolites after incorporation of ^13^C-glucose. Bar at mean ± SD, *p* values from one-way ANOVA. **d.** Heatmaps of labelling extent of glycolysis and TCA metabolites after ^13^C-glucose labelling. **e.** Mean labelling extent of TCA associated metabolites after incorporation of ^13^C-glutamine. Bar at mean ± SD, *p* values from one**-**way ANOVA with Dunnett’s correction. Heatmaps of labelling extent of TCA metabolites (**f**) and pyrimidine intermediates (**g**) after ^13^C-glutamine labelling. **h.** Diagram showing metabolic pathways, including carbon sources and inhibitors to test ATP production. **i.** Quantification of basal ATP, normalized to parental. Bars are at mean ± SD of three independent experiments. *p* values from one-way ANOVA with Tukey correction. **j.** Quantification ATP after 1 h, treatment as indicated, normalised to (**i**). **k.** Mitochondrial oxygen consumption rate (mOCR) of parental and Δ*ctr1* parasites, untreated or treated with CuSO_4_, via Seahorse assay. **l.** Metabolic map showing a shift towards a more glycolytic state. Points at mean ± SD. Quantification of basal mOCR (**m**), maximal mOCR (**n**) and basal extracellular acidification rate (ECAR) (**o**). Bars at mean ± SD of three independent experiments. *p* value from one-way ANOVA with Tukey’s correction. **p.** In gel complex IV activity assay of parental and Δ*ctr1* parasites, untreated or treated with CuSO_4_ for 24 h, demonstrating loss of complex IV activity in Δ*ctr1* parasites. TOM40 used as a loading control. Image representative of three independent experiments. **q.** In gel complex IV activity assay of parental and Δ*ctr1* parasites, copper treated as indicated. TOM40 included as loading control.

The conversion of dihydroorotate to orotate feeds electrons to complex III of the ETC via ubiquinone^48^ and inhibition of complex III has been associated with accumulation of pyrimidine intermediates^49,50^. To examine the role of complex III indirectly, we tested the sensitivity of Δ*ctr1* parasites to the apicomplexan-specific complex III inhibitor atovaquone^51^. After normalisation for growth, we found Δ*ctr1* parasites are significantly (*p* < 0.0001, extra sum of squares F test) less sensitive to atovaquone (Atq), (parental EC_50_ = 57 nM, Δ*ctr1* EC_50_ = 180 nM), indicating that disruption of copper transport leads to changes in the ETC function (**Fig. 3b**). To confirm that this phenotype reflects metabolic changes rather than non-specific growth effects, we assessed the sensitivity of *Δctr1* to the anti-parasitic drug dihydroartemisinin (DHA) and a small but non-significant (unpaired Welch’s *t* test) difference in sensitivity between Δ*ctr1* and parental parasites was observed (**Fig. S2h**). Further, copper depletion by growth in BCS-treated cells led to significantly (*p* = 0.016, one-way ANOVA) decreased sensitivity to atovaquone, which was reversed by addition of copper (**Fig. S2i**), but no change in sensitivity to pyrimethamine (**Fig. S2j**). In agreement, an active ETC and TCA cycle have been shown to be required for aspartate synthesis^52,53^ and aspartate levels are also significantly reduced in the Δ*ctr1* parasites **(Fig. 3a** and **S2e)**.

Beyond pyrimidine biosynthesis, we also saw significant changes in the abundance of metabolites of the TCA cycle (**Fig. 3a**), including decreased abundance of citrate and oxaloacetate and accumulation of malate and succinate. We interpret this pattern in light of the loss of complex IV activity in Δctr1 parasites (see below): with electron flow through the ETC blocked, we would expect a back-up of electrons through complex III and the ubiquinone pool, in turn inhibiting succinate dehydrogenase (complex II) and preventing oxidation of succinate to fumarate, consistent with the succinate accumulation we observe. Malate accumulation may arise via non-cyclic routes, such as partial reductive flux through fumarase or malic enzyme activity, that do not require continuous oxidative turnover of the cycle. By contrast, the oxidative segment of the TCA cycle (citrate to α-ketoglutarate) depends on continuous NAD+regeneration, which becomes limiting once the ETC is blocked. We suggest this explains the depletion of citrate and oxaloacetate in a TCA cycle that can no longer complete a full oxidative turn. As aspartate is maintained in near-equilibrium with oxaloacetate via aspartate aminotransferase, its depletion likely reflects this drop in oxaloacetate rather than a pyrimidine-pathway-specific effect. Similar changes in metabolites have previously been associated with increased glycolysis due to an increased requirement for NAD regeneration^54^.

Steady state metabolomics cannot illuminate flux through pathways. To investigate this, we labelled intracellular parasites for 4 h using both ^13^C-labelled glucose or glutamine, as previously described^55^. PCA analysis demonstrated good agreement between replicates (**Fig. S3a** and **S3b**). To determine overall incorporation of carbon sources by the parasites, we calculated the total fraction of labelling across all identified metabolites. Upon labelling with ^13^C-glucose we saw no significant change between conditions (**Fig. S3c**). However, when we examined only incorporation into intermediates of glycolysis or the TCA cycle, we saw significantly (one-way ANOVA with Dunnett’s correction) increased incorporation into glycolysis-intermediates, with decreased incorporation in the TCA cycle (**Fig. 3c, d** and **S3d**), both of which were rescued by addition of copper. *Toxoplasma* also incorporates carbon from glutamine into the TCA cycle^56^. In the absence of *ctr1* we see increased incorporation of glutamine-derived carbon into all detected metabolites (**Fig. S3e**) and into TCA cycle intermediates (**Fig. 3e** and **f**), potentially linked to the diversion of glucose-derived carbon away from the TCA cycle.

Given the changes in pyrimidine biosynthesis seen above, we also examined ^13^C-glutamine-labelling into pyrimidine intermediates. Incorporation of glutamine-derived carbon into pyrimidine intermediates was increased in the Δ*ctr1* parasites, however we did not detect changes in labelling of pyrimidines (e.g. UMP), suggesting inhibition of the pyrimidine biosynthesis pathway (**Fig. 3g**), in agreement with the results from the untargeted metabolomics.

We believe that Δ*ctr1* parasites are prioritising glucose for energy generation via glycolysis and funnelling glucose-derived carbon to lactate, which is secreted from the parasites. This increases the role of glutamine in macromolecular biosynthesis, explaining the increased incorporation of glutamine-derived carbon, both in the TCA cycle and across all detected metabolites.

As we simultaneously performed the labelled metabolomics with Δ*ctr1*+Cu parasites, we wanted to assess how the addition of copper for 24 h impacted metabolic flux. By PCA analysis, the Δ*ctr1*+Cu parasites were metabolically distinct from both the Δ*ctr1,* and the parental parasite lines (**Fig. S3a** and **S3b**). However, many of the changes in total labelling extent (**Fig. S3c** and **S3e**) and central carbon metabolism (**Fig. S3d** and **S3f)** appeared fully rescued by exogenous copper addition. To examine rescue metabolite labelling between the Δ*ctr1* and Δ*ctr1*+Cu conditions. Using this measure, systematically over all detected metabolites, we quantified the percentage change in addition of copper led to rescue of approx. 45% of metabolites in both glucose (**Fig. S3g**) and glutamine (**Fig. S3h**) labelling conditions, with a similar proportion of unchanged between Δ*ctr1* and Δ*ctr1*+Cu conditions. This indicates that while 24 h of excess copper is sufficient to rescue many of the metabolic changes in central carbon metabolism, some impacts of copper deficiency persist beyond 24 h.

These results demonstrate that, in the absence of copper import, *Toxoplasma* significantly rewires cellular metabolism which can be rescued by addition of excess copper.

### Copper import is required for ATP generation by the ETC

Deletion of *ctr1* led to significant metabolic changes in the parasite. To determine how these changes impacted energy production, we quantified ATP levels upon inhibition of key metabolic pathways, as previously described^24^ (**Fig. 3h**). Prior to incubation, ATP levels in the Δ*ctr1* parasite line were less than half (44%) (*p* = 0.0089, one-way ANOVA) of the parental line or when copper supplemented (**Fig. 3i**), consistent with less efficient energy generation through glycolysis. After 1 h of incubation in the presence of defined carbon sources and/or inhibitors, parental parasites generated ATP via either glycolysis (in the presence of glucose) or the TCA cycle and electron transport chain (ETC) (following incubation with glutamine) but failed to produce ATP when both pathways were inhibited (with 2-DG and oligomycin, respectively) as expected. In contrast, although Δ*ctr1* parasites produced ATP from glucose via glycolysis, no ATP production was observed following incubation with glutamine, indicating a complete loss of energy generation through the TCA cycle/ETC, a phenotype rescued by addition of copper (**Fig. 3j**).

The only known cuproenzyme in *Toxoplasma,* cytochrome c oxidase, is essential in mitochondrial respiration^25^. To determine the effect of deleting *ctr1* on parasite mitochondrial function, we assessed respiration by Seahorse assay^57^ by quantifying oxygen consumption and extracellular acidification, a proxy for glycolysis^58^. Surprisingly, we saw that in the absence of *ctr1*, there is almost no detectable mitochondrial oxygen consumption (**Fig. 3k**) with a significant decrease in both basal (**Fig. 3m**) and maximal mOCR (**Fig. 3n**) (*p* = 0.0008, and *p* = 0.0038, respectively, one-way ANOVA with Tukey’s correction). This is accompanied by a significant (*p* = 0.016, one-way ANOVA with Tukey’s correction) increase in extracellular acidification (**Fig. 3o**), suggesting an almost complete switch to a glycolytic metabolism (**Fig. 3l**). However, these metabolic effects can be fully rescued by growth of the Δ*ctr1* for 24 h in the presence of 20 µM exogenous copper (**Fig. 3k, l, m, n** and **o**). Quantification of parasite metabolism in this way must occur in extracellular parasites to avoid host cell responses. To ensure the Δ*ctr1* parasites were viable extracellularly, we quantified extracellular survival by plaque assay and found no change in survival between parental and Δ*ctr1* parasites at any timepoint tested (**Fig. S3i**), demonstrating that *ctr1* expression is not required extracellularly. Given the importance of glucose in energy generation upon deletion of ctr1, we determined if the rate of glucose uptake was changed in the absence of *ctr1*. Parasites were treated with the fluorescent glucose analogue, 2-NBDG^58^ for 30 min and levels of 2-NBDG quantified by flow cytometry. To confirm specificity, we blocked uptake using unlabelled glucose. Interestingly, despite the presumed importance of glucose uptake to Δ*ctr1* parasites, we see no change in the rate of uptake of 2-NBDG in extracellular parasites (**Fig. S3j**). This fits with our previous work showing that although glycolysis can become more fitness conferring upon disruption of mitochondrial respiration, the rate of glucose uptake does not appear to increase^58^.

Given the severe mitochondrial phenotype, we examined mitochondrial morphology by staining with the outer membrane marker TOM40^59^. However, no significant changes in the normal, loop-shaped mitochondrial morphology were seen in the Δ*ctr1* parasites (**Fig. S3k**) and mitochondrial membrane potential appeared to be maintained, based on MitoTracker staining in live parasites (**Fig. S3l**).

Our Seahorse assay indicates that Δ*ctr1* parasites appear to exhibit no oxygen consumption **(Fig. 3k)**. Oxygen is the terminal acceptor for electrons generated by the electron transport chain, specifically through the action of cytochrome c oxidase (complex IV) which transfers electrons from cytochrome c to oxygen. We hypothesise that in absence of Ctr1, copper is not delivered to cytochrome c, and complex IV is not active. To test this, we performed an in-gel complex IV activity test where reduction of cytochrome c is detected directly through the oxidation and deposition of diaminobenzidine (DAB)^60^. We found no detectable complex IV activity in the Δ*ctr1* strain, which was fully rescued by 24 h of exogenous copper supplementation (**Fig. 3p**), indicating copper uptake is essential for *Toxoplasma* complex IV activity.

The possibility of rescue of complex IV activity by copper supplementation in the absence of *ctr1* allowed us to determine the kinetics of this rescue. Δ*ctr1* parasites were incubated with copper for the indicated time and complex IV activity was subsequently assessed. Interestingly, although full rescue required 24 h of incubation, detectable complex IV activity appeared as early as 4 h post-incubation, with faint activity bands observed at 1 – 2 h post incubation. These observations suggest that the assembly machinery is pre-established, and that copper availability is the limiting factor for complex IV activation (**Fig. 3q**).

Overall, loss of *ctr1* leads to significant and widespread changes to parasite metabolism, including a switch in energy generation via glycolysis and changes in metabolic pathways which rely on the ETC. These changes can be largely and rapidly rescued by addition of exogenous copper.

### *ctr2* appears dispensable for tachyzoites lytic cycle

Having characterized Ctr1 as being the major mediator of copper uptake during the tachyzoite stage, we then turned to characterization of *ctr2*. *ctr2* seemed a likely candidate for the secondary pathway of copper import that allowed the phenotypic recovery of the Δ*ctr1* parasites in the setting of excess copper. Ctr2’s positive fitness score (2.31) suggested it is not essential for the tachyzoite life cycle and, like *ctr1*, *ctr2* appeared to be upregulated in bradyzoites^61^. Given this possibility, we utilized the CRISPR/Cas9 system to generate a *ctr2* knockout (Δ*ctr2*) in a type II strain (Prugniaud, Pru) which effectively encysts *in vivo* and *in vitro*^62^ where we confirmed gene disruption by integration PCR (**Fig. S4a, b**). We complemented the knockout line by ectopically expressing a C-terminally HA-tagged copy of *ctr2,* driven by its endogenous promotor (Δ*ctr2*:Ctr2-HA). We confirmed knockout and complementation of *ctr2* by qRT-PCR. Disruption of ctr2 did not impact the expression of *ctr1* or *ctr3* (**Fig. S4c**). Consistent with its positive fitness score, a plaque assay showed no significant differences in plaque number upon *ctr2* deletion (**Fig. 4a, b**) and quantification of parasites/vacuole at 24 h demonstrated no change in parasite replication (**Fig. 4c**).

**Figure 4.**
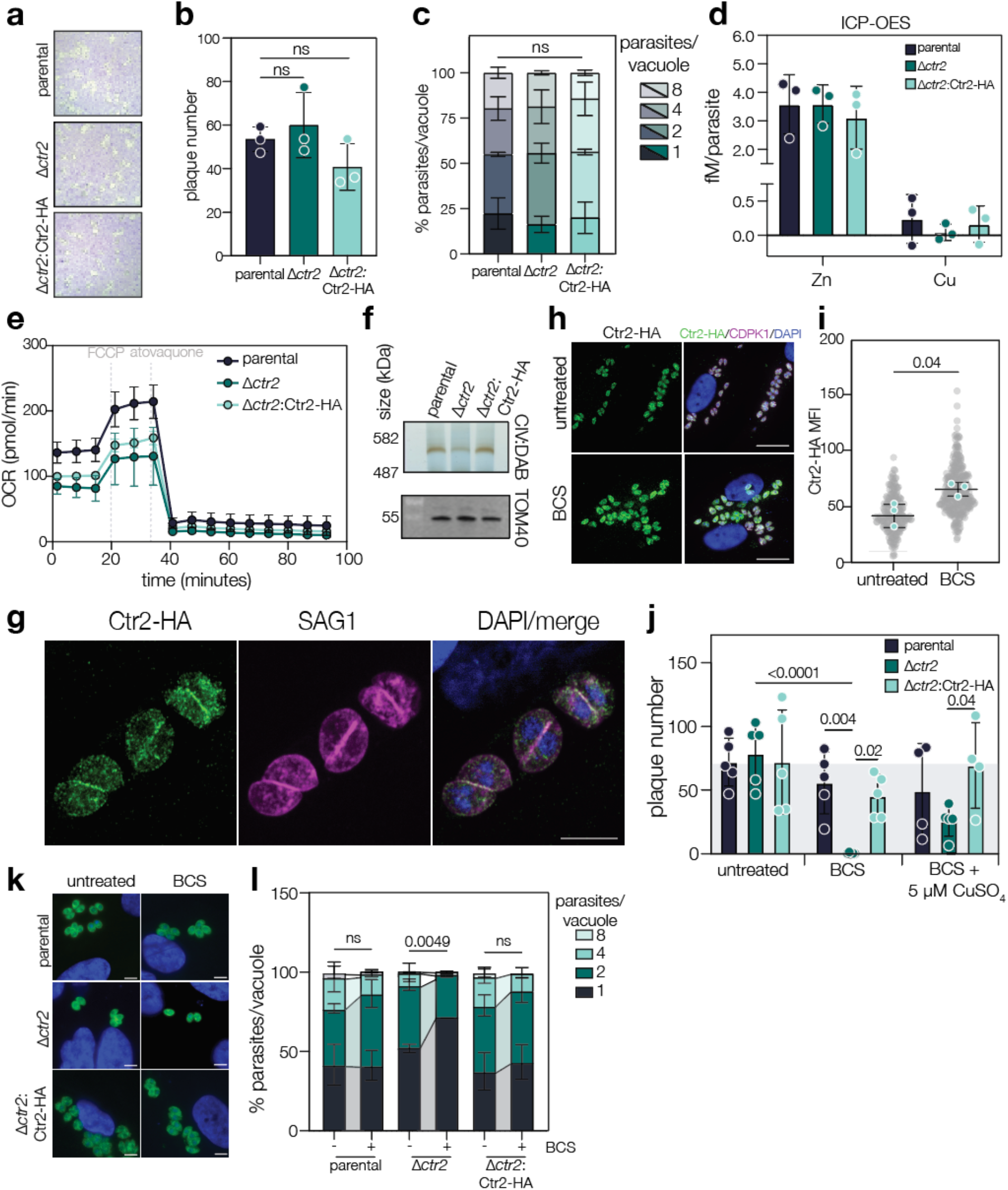
Ctr2 has a mild phenotype in tachyzoites, but is important in low copper. **a.** Plaque assay of parental, Δ*ctr2* and Δ*ctr2*:Ctr2-HA parasite lines. **b.** Quantification of plaque number. Bars are at mean ± SD*. p* values are from one-way ANOVA with Dunnett’s correction. **c.** Quantification of percentage of parasites/vacuole in parental, Δ*ctr2* and Δ*ctr2*:Ctr2-HA parasite lines parasites after 24 h of infection. 100 vacuoles for each condition were counted. Bars at mean ± SD of three independent experiments. *p* values are from two-way ANOVA with Dunnett’s correction **d.** ICP-OES from indicated parasite lines, bars at mean ± SD of three independent experiments, *p* values are from two-way ANOVA with Dunnett’s correction. **e.** Mitochondrial oxygen consumption rate (mOCR) of parental, Δ*ctr2 and Δctr2:*Ctr2-HA parasites via Seahorse assay. **f.** Complex IV activity assay of parental, Δ*ctr2* and Δ*ctr2*:Ctr2-HA parasites. TOM40 used as a loading control. **g.** Immunofluorescence assay of Ctr2-HA at 24 h post infection showing intracellular vesicular localisation. Scale bar 5 µm. **h.** Immunofluorescence assay of Ctr2-HA at 24 h post infection showing increased expression under copper depletion. Scale bar 20 µm. **i.** Quantification of mean fluorescence intensity (MFI) of Ctr2-HA. Bars are at mean ± SD of three independent experiments. *p* values from unpaired two-tailed *t* test with Welch’s correction. **j.** Plaque numbers performed under untreated, BCS (copper depleted) or BCS + Cu conditions. Bars at mean ± SD, *p* values are from one-way ANOVA with Tukey’s correction. **k.** Representative images of parental, Δ*ctr2* or Δ*ctr2*:Ctr2-HA parasite lines in untreated or BCS copper depleted conditions. Scale bar 20 µm. **l.** Quantification of percentage of parasites/vacuole in indicated parasite lines under untreated or BCS treated conditions, at 24 h post infection. Bars at mean ± SD of three independent replicates, p value from two way ANOVA with Sidak correction.

To determine if *ctr2* was important in copper uptake, we then quantified copper levels in the parasites using ICP-OES. We saw no change in parasite-associated copper or zinc upon deletion of *ctr2* or in the complemented line, suggesting that *ctr2* is not required for copper uptake in *T. gondii,* in the presence of *ctr1* (**Fig. 4d**).

Our data show that *ctr1* is required for mitochondrial respiration. To determine if there are changes in mitochondrial respiration upon deletion of *ctr2*, we performed a Seahorse assay (**Fig. 4e**), as above. Our results show that Δ*ctr*2 parasites have a significantly (*p* = 0.01, one-way ANOVA with Tukey’s correction) reduced basal oxygen consumption rate, when compared to the parental line (**Fig. S4d**). We also found a significant decrease (*p* = 0.004, one-way ANOVA with Tukey’s correction) in maximal (**Fig. S4e**) oxygen consumption in the Δ*ctr*2 parasites, although changes in OCR were not restored in the complemented line. However, we saw no significant change in the ECAR in any of the parasite lines (**Fig. S4f**), leading to only a small shift in the energetic phenotype of the parasite (**Fig. S4g**). To determine if complex IV activity was mediating this change, weanalysed complex IV activity as described above. In contrast to Δ*ctr1* parasites, where complex IV activity was totally abolished, our results show that complex IV oxidation activity was only slightly reduced when compared to the parental line and fully rescued in the complemented line (Δ*ctr2*:Ctr2-HA) (**Fig. 4f**).

To determine the localization of Ctr2-HA, we performed an immunofluorescence assay against the HA epitope and found Ctr2-HA localized to numerous internal vesicular structures within the parasite, with limited staining at the plasma membrane (**Fig. 4g**). We quantified Ctr2-HA abundance and found a significant (*p* = 0.04, unpaired *t* test) increase of Ctr2-HA upon copper depletion (**Fig. 4h, i**). We also saw a statistically significant increase in peripheral localization of Ctr2-HA under copper depletion (*p* < 0.0001, two-way ANOVA, Fisher LSD test) (**Fig. S4i, j**). These results suggest a role of Ctr2 in low copper environments.

### Ctr2 is essential under low copper

As we had already determined that *ctr2* was not fitness conferring in nutritionally replete conditions, we next tested the requirement for *ctr2* in copper deficiency using BCS-treated cells. Upon quantification of plaque numbers, we found that Δ*ctr2* formed almost no visible plaques in BCS-treated host cells (*p* < 0.0001, two-way ANOVA with Tukey correction). This growth defect was rescued either by copper supplementation, or by Ctr2 expression in the Δ*ctr2*:Ctr2-HA parasite line (**Fig. 4j**), demonstrating copper-specificity. To confirm this result, we quantified number of parasites/vacuole. At 24 h post infection, we saw significantly more (*p* = 0.0049, two-way ANOVA with Sidak correction) 1-parasite vacuoles in BCS-treated cells, compared to untreated (**Fig. 4k**, **l**), but only in the Δ*ctr2* parasite line. These results demonstrate that loss of *ctr2* leads to increased sensitivity to copper depletion.

These data demonstrate that *ctr2*, although largely dispensable under *in vitro* lab conditions, has a role in tachyzoite survival in low copper environments. Its increased expression and change in localization also support our hypothesis that *ctr2* can compensate for loss of *ctr1*, although with reduced effectiveness.

### Copper transport is not required for *in vitro* differentiation

Both *ctr1* and *ctr2* are highly expressed in bradyzoites, suggesting a potential role in this lifecycle. We first examined the localisation of Ctr1-mNG in parasites grown under alkaline stress conditions for 7 days and confirmed that the localisation is maintained at the parasite membrane (**Fig. 5a**). To determine if copper import is required for bradyzoite conversion, we infected cells with parental and Δ*ctr1* parasites for 24 h then placed in alkaline stress conditions, with or without additional copper. After 7 days we fixed and stained using *Dolichos biflorus* lectin (DBL), which recognizes sugar modifications in the cyst wall proteins^63^. We saw no significant changes in the % DBL-positive vacuoles (**Fig. 5b**), suggesting that copper import via Ctr1 is not essential for bradyzoite conversion. We also quantified cyst area and found that DBL-positive structures were significantly (*p* = 0.0001, one-way ANOVA, Tukey’s comparison) smaller in the Δ*ctr1* parasites, similar to the tachyzoite growth defect, suggesting that copper is important for bradyzoite replication (**Fig. 5c, d**). In contrast to our previous results, addition of copper did not fully rescue this phenotype (**Fig. 5c, d**).

**Figure 5.**
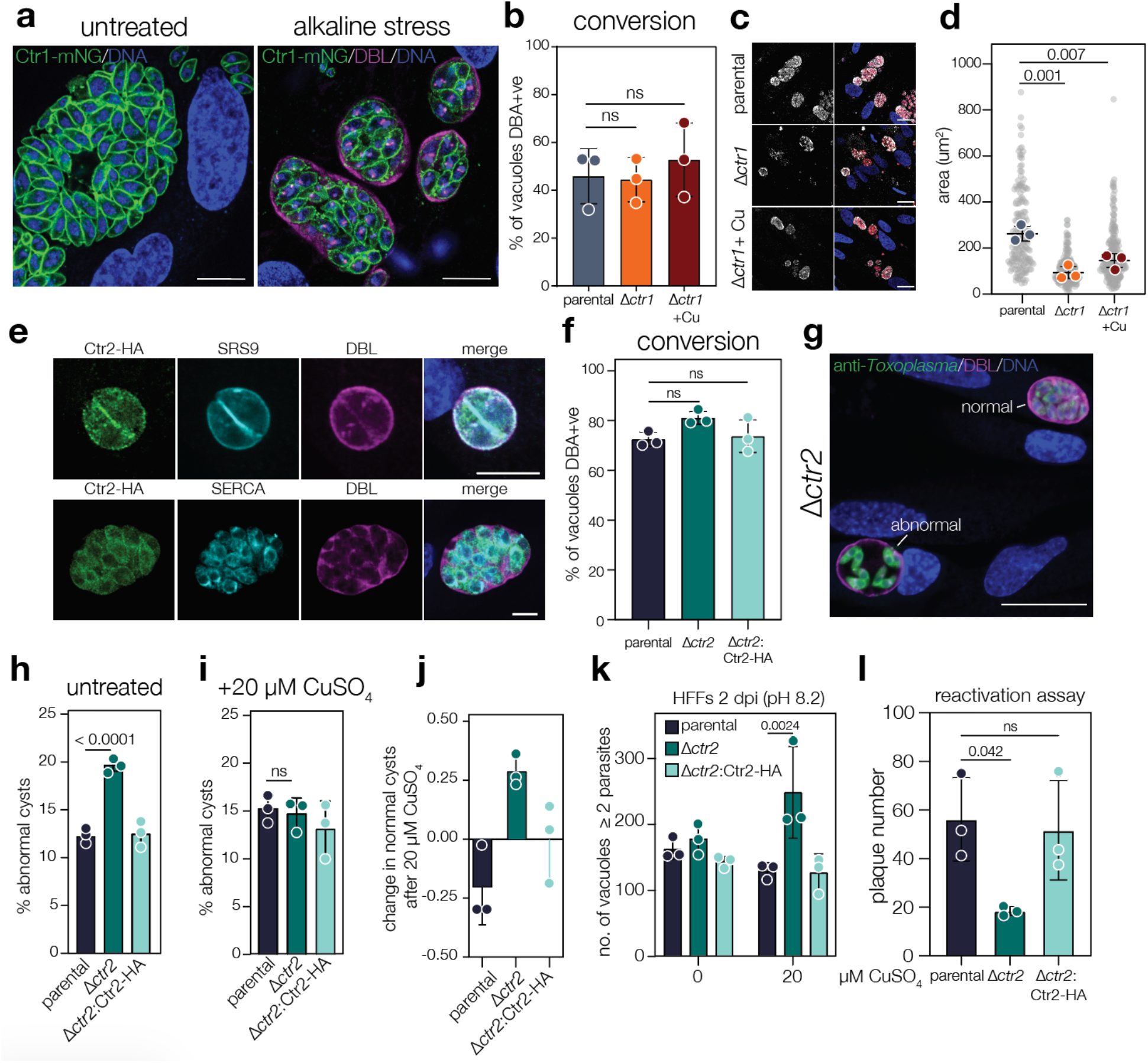
Role of copper transporters in bradyzoites. **a.** Ctr1-mNG is peripherally localised in untreated and in DBA-positive vacuoles at seven days of alkaline stress. Scale bar 10 µm. **b.** Quantification of % of DBA-positive vacuoles by parental or Δ*ctr1* parasites and Δ*ctr1* parasites with copper supplementation. Bars are at mean ± SD, *p* value from two-way ANOVA with Tukey’s correction **c.** Immunofluorescence of DBA-positive vacuoles formed by parental or Δ*ctr1* parasites with or without copper supplementation upon seven days of alkaline stress. Scale bar 20 µm. **d.** Quantification of DBA-positive vacuolar area. Coloured points are the mean of three independent experiments, bar are at mean ± SD. *p* values are from two-way ANOVA with Sidak correction. **e.** Immunofluorescence assay of Ctr2-HA upon two days of *in vitro* bradyzoites differentiation by alkaline stress. Scale bars 5 μm. **f.** % of vacuoles with DBL staining after 2 days alkaline stress. Bars are at mean ± SD, *p* value from one-way ANOVA with Dunnett’s correction from three independent experiments. **g.** Immunofluorescence assay of Δ*ctr2* after differentiation, showing the formation of abnormal cysts. Scale bar 20 µM. **h.** Quantification of abnormal cysts. Bars are at mean ± SD, *p* value from two-way ANOVA with Tukey’s correction from three independent experiments. **i.** Quantification of abnormal cysts after 20 µM CuSO_4_ treatment. Bar at mean ± SD, *p* value from two-way ANOVA with Tukey’s correction from three independent experiments. **j.** Change in normal cysts across genotypes, after addition of 20 µM CuSO_4_, only in the Δ*ctr2* parasite line does addition of copper prevent abnormal cysts. **k**. Quantification of number of vacuoles (under alkaline stress) after copper supplementation. Bars are at mean ± SD, *p* value from two-way ANOVA with Tukey’s correction from three independent experiments. **l.** Quantification of plaque number after bradyzoites reactivation. Bars are at mean ± SD, *p* value from one-way ANOVA with Dunnett’s correction from three independent experiments.

We also analysed Ctr2 localisation upon growth under alkaline stress, and see no significant changes, with the majority of signal distributed in the cytosol with some peripheral localization, which overlapped with SRS9 (**Fig. 5e**). To determine if *ctr2* has a role in bradyzoites formation, we quantified vacuole differentiation (defined by positive DBL staining) using an automated pipeline after two days of alkaline stress, as previously described^64^. Using this method, we saw no statistically significant difference in percentage of DBL-positive cysts after conversion in alkaline media (**Fig. 5f**). Looking more closely, we saw a slight decrease in the total number of vacuoles or cysts in the absence of *ctr2*, although this was not significant (**Fig. S4k, l**). We also determined the rate of conversion after growth in neurons, representing more physiological conditions. However, there was no significant change in conversion rate in these conditions (**Fig. S4m**), suggesting *ctr2* is not essential for parasite differentiation.

However, we noticed that many cysts exhibited abnormal morphology, with fewer and more dispersed parasites (**Fig. 5g**). We quantified this phenotype using a custom machine-learning phenotype classification pipeline and found that deletion of *ctr2* led to a significant increase in abnormal cysts (**Fig. 5h**) (*p* < 0.0001, two-way ANOVA with Tukey’s correction), which was rescued in the complemented strain. Interestingly, this increase in abnormal cysts was also rescued by incubation with 20 µM CuSO_4_ (**Fig. 5i, j**) suggesting that the formation of abnormal cysts is a result of copper starvation in the Δ*ctr2* parasites. Supporting this, we find a significantly (*p* = 0.0024, two-way ANOVA with Tukey’s correction) higher number of vacuoles with > 2 parasites in the Δ*ctr2* parasite line upon copper supplementation (**Fig. 5k**), although this is not seen in parental or complemented strains, suggesting that exogenous copper can promote growth in the absence of *ctr2*.

To determine if loss of *ctr2* and accumulation of abnormal cysts affects parasite survival, we tested the viability of Δ*ctr2* bradyzoites through a reactivation assay. In this assay, fibroblasts were seeded with 250 parasites of each strain, followed by encystment conditions for 6 days, then a return to lytic conditions for 14 days, after which cultures were fixed, stained, and plaques counted. The Δ*ctr2* parasites formed significantly (*p* = 0.042, one-way ANOVA with Dunnett’s correction) fewer plaques than the parental or complemented parasite lines (**Fig. 5l**), suggesting that abnormal cysts have fewer viable parasites.

These data show that copper import is not required for differentiation, however *ctr2* plays a role in parasite fitness during stage conversion and cyst maintenance.

### Δ*ctr2* parasites are cleared during acute infection *in vivo*

Given that, *in vitro*, Δ*ctr2* parasites had a normal lytic cycle but formed abnormal cysts, we sought to test the requirement for *ctr2* during *in vivo* encystment. Mice were infected intraperitoneally with parental, Δ*ctr2* and Δ*ctr2*:Ctr2-HA strains and weight, as a surrogate for infection severity, was monitored for 3 weeks. For the first 10 days, no significant differences were observed in weight loss experienced by mice infected with each strain (**Fig. 6a**). However, from 12 days post infection (dpi), mice infected with Δ*ctr2* or Δ*ctr2*:Ctr2-HA parasites began to regain weight and exhibit significantly less weight loss relative to mice infected with the parental strain (two-way ANOVA, Dunnett’s multiple comparison), potentially suggesting a milder infection.

**Figure 6.**
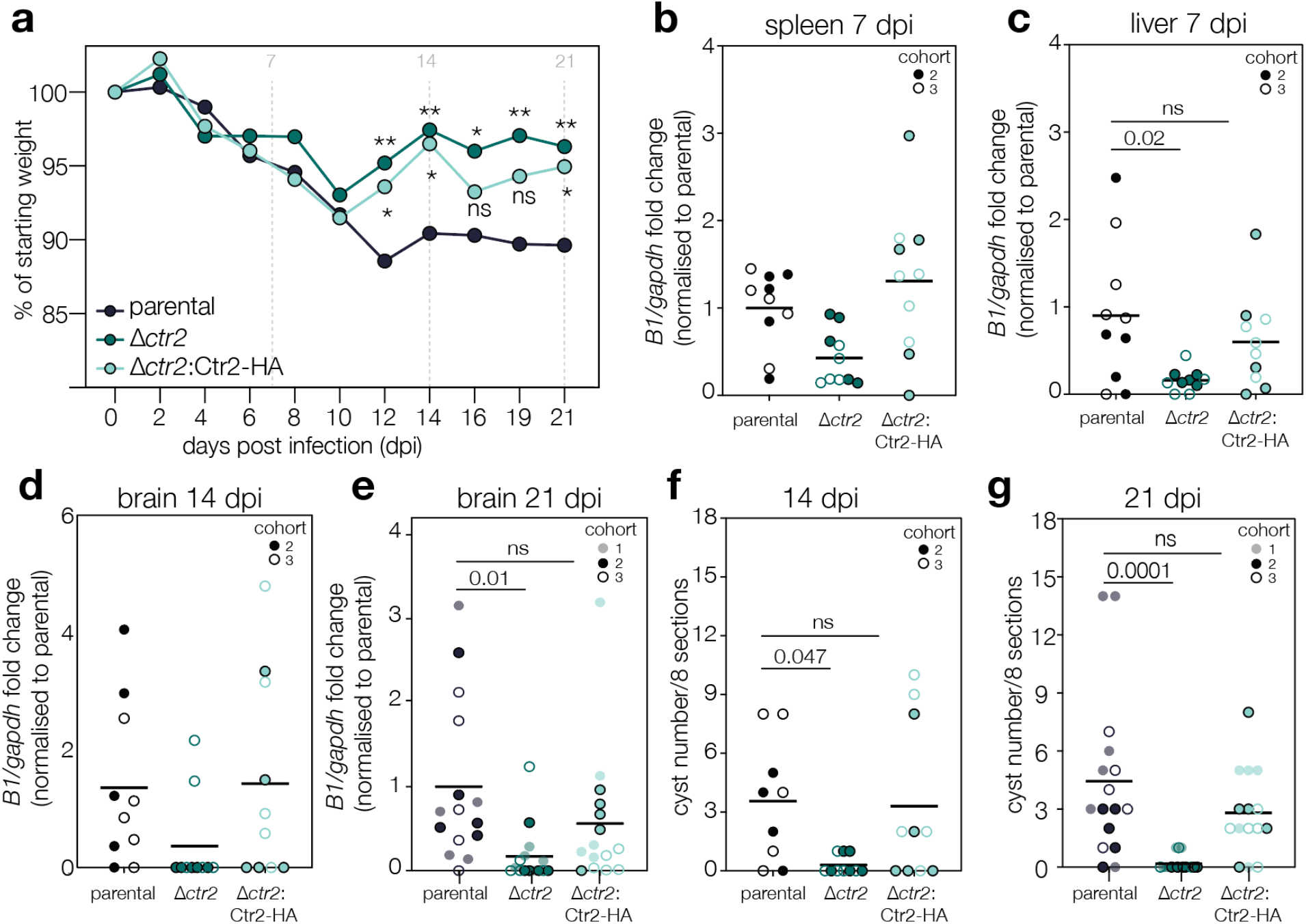
Role of *ctr2 in vivo*. **a.** Mice weight during acute infection, * *p* < 0.05, ** *p* < 0.005, significance from two-way ANOVA with Dunnett’s correction. Parasite burden by qPCR in spleen **(b)** or liver **(c)** at 7 dpi., normalised to host GAPDH as housekeeping control. . *p* values from two-way ANOVA with Dunnett’s correction, Parasite burden by qPCR from the brain at 14 (**d**) or 21 (**e**) dpi, *p* value from two-way ANOVA with Dunnett’s correction. Quantification of cyst number in brain sections at 14 (**f**) or 21 (**g**) dpi. *p* value from two-way ANOVA with Tukey’s correction. For all graphs, each point represents an individual mouse, cohort indicated by symbol, lines at grand mean of all cohorts.

To investigate dissemination during infection, spleens and livers of two independent cohorts of mice (5 mice/condition/cohort) were harvested at 7 dpi and parasite burden evaluated by qPCR for the *Toxoplasma*-specific gene B1. Consistent with our prior work^65^, although all strains were detectable in the spleen and liver at 7 dpi (**Fig. 6b** and **c)**, we saw a decrease (although not significant) in Δ*ctr2* parasite burden in the spleen and a significant (*p* = 0.02, two-way ANOVA, Dunnett’s correction) decrease in parasites in the liver. To determine if Δ*ctr2* parasites were able to reach the brain, we quantified parasite DNA from harvested brains at 14 and 21 dpi (**Fig. 6d, e**). At 14 dpi we occasionally detected parasite DNA in brains of infected mice, however by 21 dpi we saw a significant (*p* = 0.01, two-way ANOVA with Tukey’s correction) decrease in detectable parasite DNA. As DNA may be present in the absence of viable parasites, and so we also quantified cysts from brain sections at both 14 and 21 dpi (**Fig. 6f, g**). At both time points we saw a significant (*p* = 0.01, two-way ANOVA with Tukey’s correction) drop in cyst counts in the Δ*ctr2* parasite line, confirming the lack of dissemination in the Δ*ctr2* parasites.

Collectively, our data show that deletion of *ctr2* results in an early survival defect *in vivo*, leading to decreased parasite dissemination and significantly decreased persistence in the brain.

## Discussion

This study provides the first analysis of copper transport and function across the asexual lifecycle of *Toxoplasma gondii*. Here we establish roles for copper transport in mitochondrial respiration and in parasite dissemination, both *in vitro* and *in vivo*.

Copper-dependent transcriptional regulation has been previously observed in diverse organisms^66–68^ and we find that both *ctr2* and *ctr3* are transcriptionally responsive to copper availability, although in opposite directions. This is not unprecedented, the iron transporters DMT1 and ferroportin are both typically regulated via the iron-regulatory protein (IRP) where high iron leads to increased ferroportin and decreased DMT1 levels^69,70^. These data contrast with the recently described *Toxoplasma* iron and zinc transporter ZFT which is only post-transcriptionally regulated by TgPro binding to *zft* mRNA^9,71^. We do not yet understand the mechanisms of copper responsiveness in *T. gondii*. Indeed, how apicomplexan parasites respond to environmental signals remains in general poorly understood. These results suggest the presence of distinct, currently uncharacterised regulatory networks for transition metal uptake. Here, we do not further address the role of Ctr3. This gene is expressed at very low levels both *in vitro* and *in vivo* and is upregulated in excess copper. In yeast, Ctr3 is redundant with Ctr1 but has distinct regulatory properties^72^. Here, we found that *ctr3* is not significantly transcriptionally regulated in the Δ*ctr1* parasite line, however it may play a role in the rescue of the Δ*ctr1* parasites under high copper. Under standard conditions, it does not appear that *ctr2* or *ctr3* are sufficient to complement for the loss of *ctr1*.

Our data show that *ctr1* is both necessary and sufficient for copper uptake in *Toxoplasma* under standard growth conditions. This conclusion is based on the ability of Ctr1 to mediate copper transport in *Xenopus* oocytes and that in its absence, parasite-associated copper becomes almost undetectable. We were able to visualise this by utilising recent advances in high-resolution, cryo-XFM, where we can localise copper to subcellular structures. Copper is seen in small, concentrated foci which frequently overlap with iron, zinc and calcium. Copper and iron uptake are tightly linked in a range of species, due to a requirement for copper-dependent iron reductases in iron uptake^73–75^. However, we did not see changes in parasite associated iron upon deletion of *ctr1,* suggesting limited crosstalk between these metal uptake systems and supporting our previous hypothesis that *Toxoplasma* does not encode iron reductases^9^. Endogenous tagging showed Ctr1 is localised to the parasite plasma membrane with a pool of protein at the Golgi, as predicted by hyperLOPIT^76^. In mammalian cells, Ctr1 localization is dynamic and is trafficked between intracellular vesicles and the plasma membrane in a copper-dependent manner^77^. This does not appear to be the case in *Toxoplasma* as Ctr1-mNG does not significantly change in localisation upon copper supplementation or depletion. As mechanisms of transporter regulation are complex, it is possible that Ctr1 activity changes in response to environmental copper concentrations.

Given the central role of Ctr1 in copper uptake, Δ*ctr1* parasites provided a tool to examine the role of copper in *Toxoplasma* biology. We found that copper is essential for complex IV activity. The lack of copper leads to a complete block of ATP generation through the ETC, inhibition of flux through the pyrimidine biosynthesis pathway, and a major metabolic shift to aerobic glycolysis. This shift resembles a Warburg-like effect, as Δ*ctr1* parasites use glycolysis to generate energy while maintaining the TCA cycle to produce intermediates required for macromolecule biosynthesis^58^. Notably, a similar metabolic phenotype has previously been described in *Toxoplasma* mutants deficient in haem-biosynthesis (CPO knockout parasites)^78^. Together with our findings, this suggests that *Toxoplasma* possesses a conserved metabolic program which allows for survival with minimal ETC activity, regardless of the specific upstream cause. Supporting this hypothesis is the surprising finding that although the mitochondrial ETC is non-functional, the mitochondrial membrane potential is preserved. Similar results have been seen in other organisms^79,80^, however not in *Toxoplasma* to our knowledge. Interestingly, this profound metabolic shift is not reflected in the transcriptome, demonstrating post-transcriptional metabolic but not transcription adaptation in the Δ*ctr1* parasites.

Copper supplementation rescues the metabolic phenotype, including the complex IV defect. After internalization, mitochondrial copper is passed through a relay of chaperones before its insertion into complex IV. Several of these key chaperones, including *cox11* (TGME49_202800), *cox17* (TGME49_240550), *cox19* (TGME49_254260) and *sco1* (TGME49_257440) are conserved in Apicomplexa, where they likely perform a similar process. We found that exogenous copper can rapidly start to rescue the complex IV defect, with activity returning within 4 h of copper supplementation. Studies of human complex IV assembly suggest that copper is inserted into cox2 (which in *T. gondii* is encoded by two genes, *cox2a* and *cox2b*) via a copper metallochaperone complex, prior to complex assembly and haemylation^81^. The rapid kinetics of complex IV activity rescue observed in *Toxoplasma* suggest that ETC subunits continue to be synthesised in the absence of copper. Similar kinetics were previously suggested in yeast^82^, although in human cells incorporation of copper into complex IV can take up to 48 h^83^. Although the structure and composition of *Toxoplasma* complex IV has been solved^19,20^, its assembly, metalation and regulation have not yet been determined. Our results suggest that synthesis of the subunits continues in the absence of assembly, and that copper internalization (including transit through the host cell) and incorporation can occur within hours.

The rapid rescue of the Δ*ctr1* line upon copper supplementation suggests that *T. gondii* possesses an alternative route for copper acquisition. Ctr2 is a candidate for this role. In both human and yeast cells, Ctr2 mobilises copper from intracellular stores and has the potential to enable copper import. Ctr2 has been localized to the endosomal system (mammalian cells) or vacuole (yeast), with limited or transient localization to the plasma membrane^84,85^. In contrast to *ctr1*, we were able to complement *ctr2* with a tagged version (under the endogenous promoter) and localise it within the parasite. Using our complemented line, we find *Toxoplasma* Ctr2 typically localises to internal vesicles, with some relocalisation to the plasma membrane upon copper depletion, where it is potentially available for copper import. Direct transport activity for Ctr2 has not been demonstrated, and further work will be required to establish whether Ctr2 mediates copper uptake at the parasite plasma membrane and/or contributes to copper mobilisation from intracellular stores.

Although deletion of *ctr2* had no significant effect on parasite growth or respiration in standard conditions, Ctr2 was required in low copper conditions, had a role in cyst maintenance, and during *in vivo* infection. These phenotypes may be mechanistically linked, as the media for the alkaline encystment model uses only 1% serum^86,87^, which is likely to have less available copper. Similarly, *in vivo*, sequestration of nutrients away from pathogens (including essential metals) is an important aspect of host defences^3,88,89^. Together, these observations suggest that Ctr2 is important in situations where copper is not freely available, and that copper availability is limited during in vivo infection, although the locations and dynamics of this are unknown.

The role of copper in bradyzoites remains unknown. Copper is not required for triggering conversion, as knockout of either transporter did not impact conversion rates, although with the caveat that the Δ*ctr1* experiments were performed in the RH strain which is defective in bradyzoite formation. Tachyzoites appear to have a robust requirement for copper in the ETC, however bradyzoites, which replicate slowly and have decreased reliance on energy generation through the ETC^90,91^ may have a reduced requirement. Loss of copper in the Δ*ctr1* line led to smaller but still apparently functional and replicating DBA-positive vacuoles, suggesting that the metabolic adaptation which sustained the tachyzoites was sufficient in bradyzoites. It is possible that there are additional roles for copper at this lifecycle stage, especially given the copper-dependent accumulation of abnormal cysts in the Δ*ctr2* parasite line, presenting the opportunity for future studies to build on these findings.

Here, we have demonstrated the importance of copper to the asexual biology of *Toxoplasma gondii* through genetic manipulation of two key copper transporters. We identify Ctr1 as the major copper importer of the parasite and show that Ctr2 is required for efficient, persistent infection. Despite these advances, fundamental questions remain regarding intracellular copper trafficking, detoxification, copper-dependent transcriptional regulation, and the precise role for Ctr2. Nonetheless, this work represents an important advance in our understanding of the metallobiology of these intracellular parasites and highlights the metabolic flexibility behind the adaptation to fluctuating host environments.

## Supplemental figures

**Figure S1.**
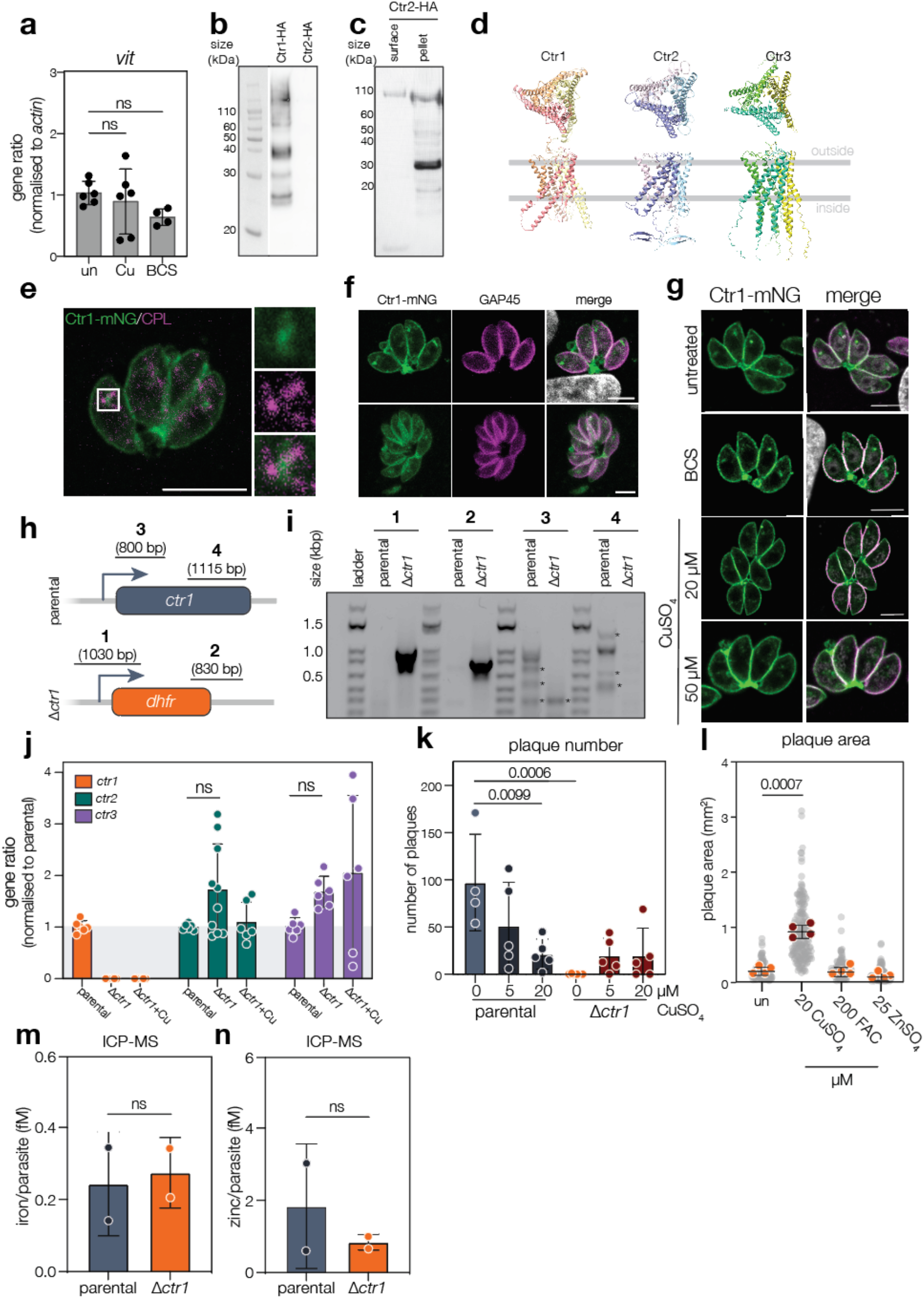
Construction and characterisation of the Δ*ctr1* parasite line. **a.** Quantification of *vit* transcripts, normalized to actin as housekeeping control, in untreated, copper treated or Cu-depleted (BCS) parasites. Bars are at mean ± SD of three independent experiments, *p* values from one-way ANOVA with Tukey’s correction. **b**. Western blot showing surface expression of Ctr1-HA (expected size 42 kDa), but not Ctr2-HA in injected *Xenopus* oocytes. **c**. Western blot showing expression of Ctr2-HA, but no membrane localisation**. d.** AlphaFold models of Ctr1, Ctr2 and Ctr3 showing predicted trimeric structure. **e.** Ctr1-mNG does not co-localise with the PLVAC marker CPL. Scale bar 5 µm. **f**. Ctr1-mNG localisation does not change based on number of parasites/vacuole. Scale bar 5 µm. **g.** Ctr1-mNG parasites were treated for 24 h, as indicated, localisation of Ctr1 did not change under these conditions. Scale bar 5 µm. **h.** Schematic summarizing the construction of the Δ*ctr1* parasite line. **i.** PCR confirmation of replacement of the coding region of ctr1 with the *dhfr* cassette. **j**. Quantification of *ctr1, ctr2 and* ctr3 transcripts, normalized to actin as housekeeping control, in parental, Δ*ctr1* and Δ*ctr1 +*copper treated parasites. Bars are at mean ± SD of indicated independent experiments, *p* values from one-way ANOVA with Tukey’s correction. **k.** Number of plaques of parental and Δ*ctr1* after addition of indicated copper concentrations. Bars are at mean ± SD of three independent experiments. *p* value from one-way ANOVA with Tukey’s correction. **l**. Plaque area after treatment of Δ*ctr1* with the indicated metals. Gray indicates all points, coloured points show mean of 4 independent experiments, lines at mean ± SD. *p* values from one-way ANOVA with Tukey’s correction. Iron (**m**) and zinc (**n**) concentrations from parental and Δ*ctr1* parasites measured by ICP-MS showing no significant difference between strains (*p* value from unpaired t test).

**Figure S2.**
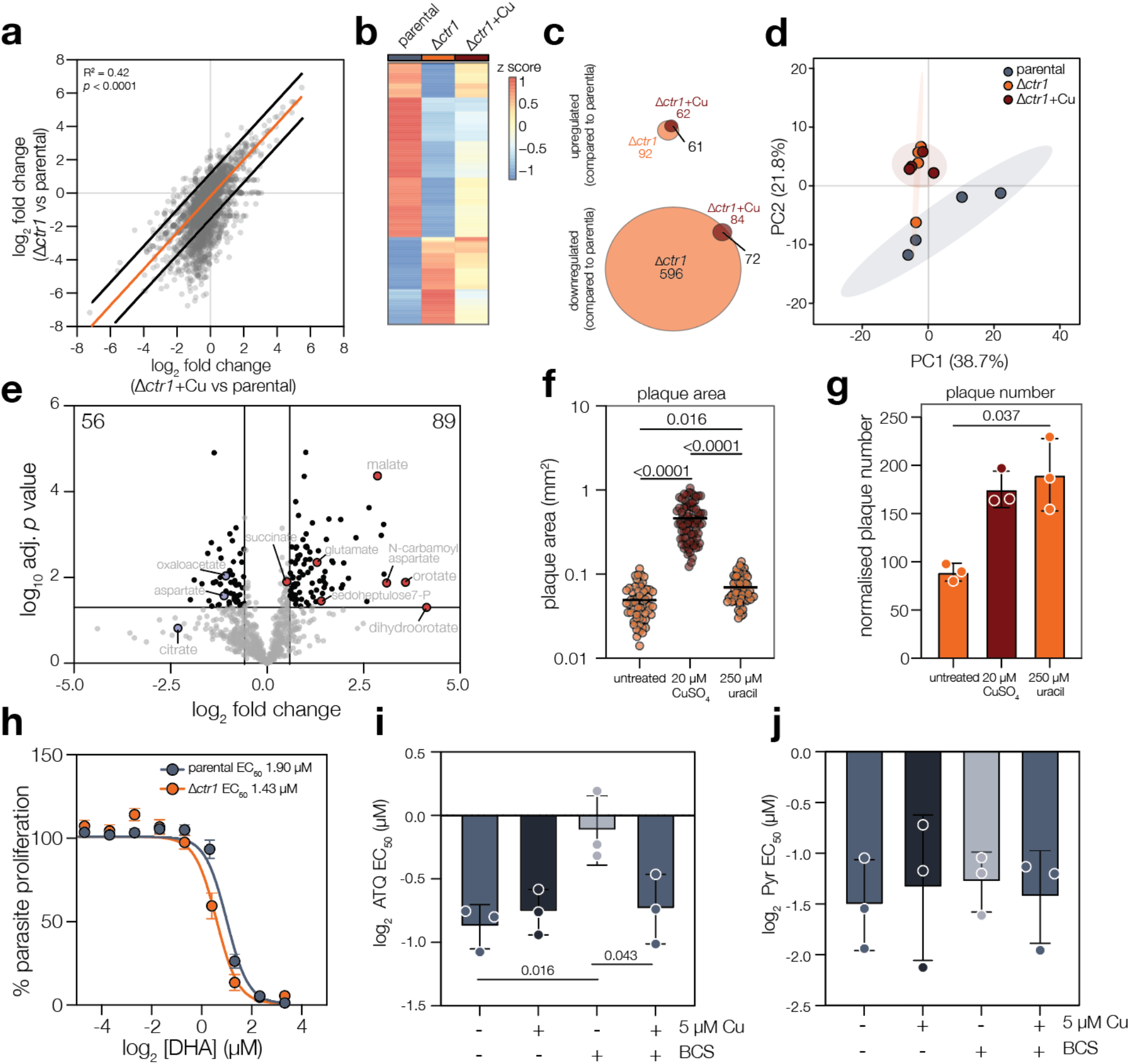
Phenotypic characterisation of Δ*ctr1* parasite line. **a.** Log_2_ fold change in transcript abundance of Δ*ctr1*/parental and Δ*ctr1*+Cu/parental. Line from linear regression. **b.** Heatmap of 150 highest DEGs between parental and Δ*ctr1* parasites, showing overall rescue of gene expression after 24 h growth with exogenous CuSO_4_. **c**. Venn diagrams of significantly (adj.p < 0.05, log_2_ fold change < -1.5 or > 1.5) regulated genes between parental and Δ*ctr1* and parental and Δ*ctr1*+Cu. The overlap (in black) indicates genes which are significantly up- or downregulated in both conditions. **d.** PCA plot showing variation between replicates for untargeted metabolomics showing untreated and copper treated conditions, ellipses represent the 95% confidence interval. **e**. Volcano plot of differential metabolite abundance with log_10_ adjusted *p*-value plotted against the log_2_ fold change (Δ*ctr1*/parental). Significant (adj. *p* < 0.05, log_2_ fold change (L_2_FC) < -0.6 or > 0.6) metabolites indicated in black. Plaque area (**f**) and normalised plaque number (**g**) of Δ*ctr1* parasites, treated as indicated, from three independent replicates. Bar at median (**f**) or mean ± SD (**g**), *p* value from one-way ANOVA with Dunnett’s correction. **h.** Fluorescent growth assay of parental and Δ*ctr1* parasites upon treatment with dihydroartemsinin (DHA). Points are mean, n = 3 ± SEM. Log_2_ EC_50_ of parasites treated as indicated to atovaquone (**i**) and pyrimethamine (**j**). Bars at mean of n = 3, ±SD. *p* values from one-way ANOVA with Dunnett’s correction.

**Figure S3.**
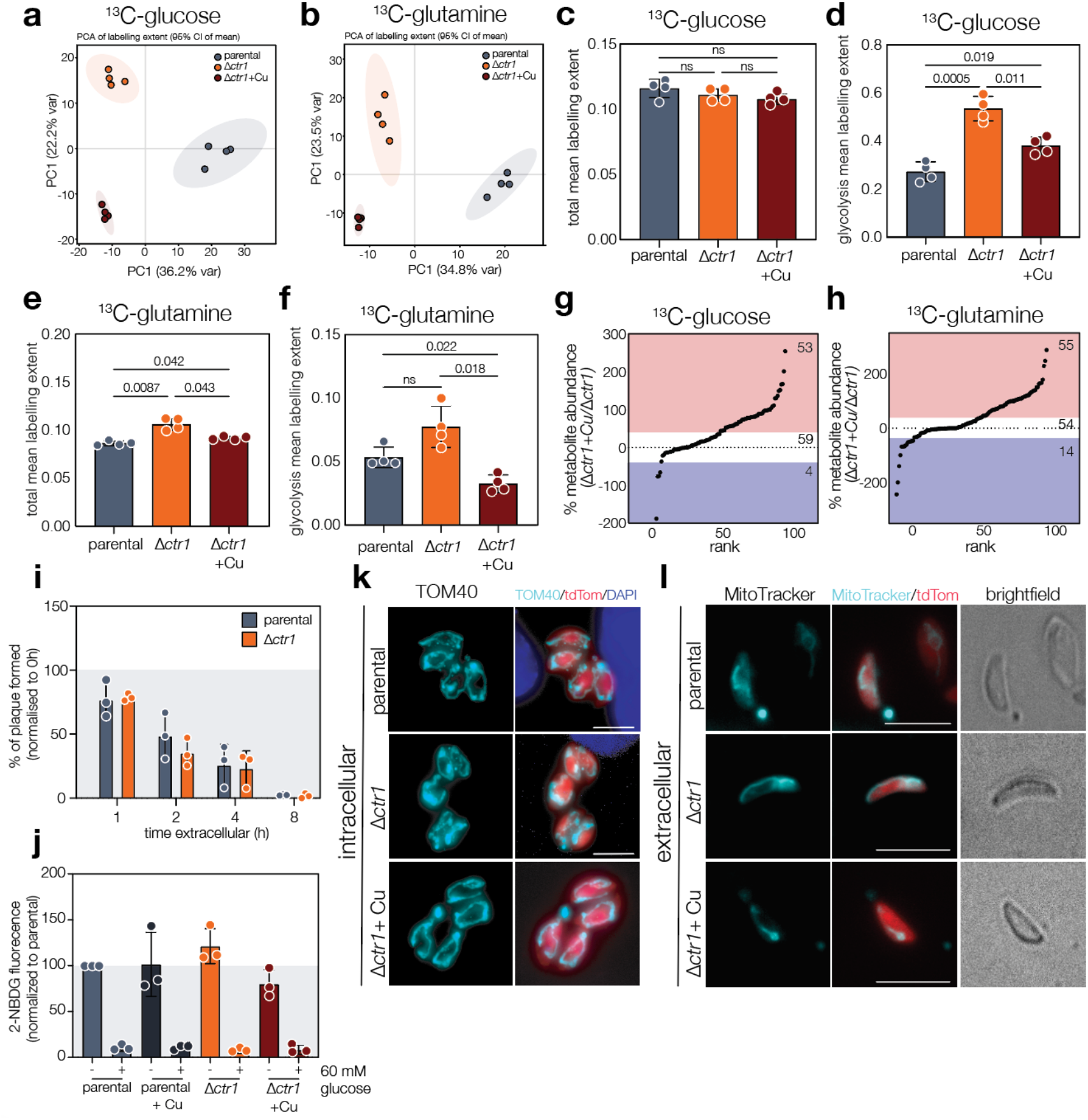
Phenotypic and metabolic characterisation of Δ*ctr1* parasite line. PCA plots of labelling extent after labelling with ^13^C-glucose (**a**) or ^13^C-glutamine (**b**), ellipses represent the 95% confidence interval. Bar graphs of average ^13^C-glucose labelling extent (LE) for all metabolites (**c**) and only those involved in glycolysis (**d**). Bar at mean ±SD, *p* values from one-way Brown-Forsythe and Welch ANOVA with Dunnett’s correction. Bar graphs of average ^13^C-glutamine LE for all metabolites (**e**) and glycolysis only (**f**) for each condition. Bar at mean ±SD, points represent replicates, *p* values from one-way Brown-Forsythe and Welch ANOVA with Dunnett’s correction. **g**. Comparison of % of rescue of metabolite labelling after ^13^C-glucose between Δ*ctr1*and Δ*ctr1*+Cu conditions. Lines at 1 SD from mean. **h**. Comparison of % of rescue of metabolite labelling after ^13^C-glutamine between Δ*ctr1*and Δ*ctr1*+Cu conditions. Lines at 1 SD from mean. i. Extracellular survival of parental and Δ*ctr1* parasites, normalized to 0h. No differences between parental and Δ*ctr1* were seen. Bars at mean ± SD of three independent experiments. j. 2-NBDG fluorescence, normalized to parental untreated. 2-NBDG was outcompeted by addition of 60 mM glucose in all strains, however no differences in 2-NBDG uptake were observed between strains. Bars are at mean ± SD of three independent experiments. k. Mitochondrial morphology examined using TOM40, no gross morphological changes could be seen in intracellular parasites. Scale bar 5 μm. **l.** Live, extracellular parental and Δ*ctr1* parasites, stained with MitoTracker, showed no gross differences in mitochondrial morphology. Scale bar 5 μm

**Figure S4.**
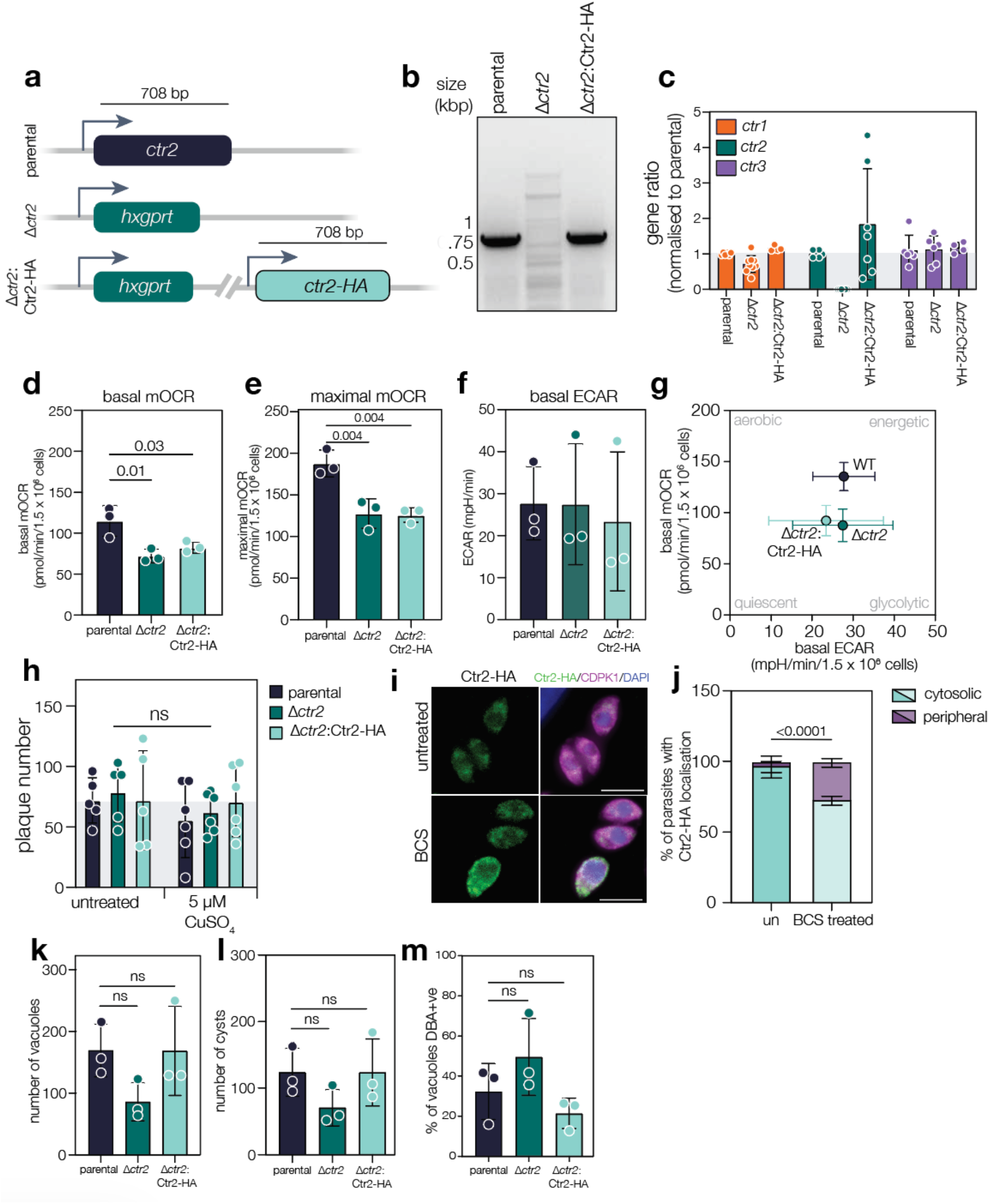
Phenotypic characterisation of the Δ*ctr2* strain. **a.** Schematic demonstrating method for strain engineering the Δ*ctr2* and Δ*ctr2*:Ctr2-HA strains. **b.** PCR showing loss of native *ctr2* coding region in the Δ*ctr2* strain. **c.** Quantification of *ctr1*, *ctr2* and *ctr3* transcripts, normalized to actin as housekeeping control, by qRT-PCR in parental, Δ*ctr2* and Δ*ctr2*:Ctr2-HA parasite lines. Bars are at mean ± SD from indicated independent experiments. *p* values from two-way ANOVA with Sidak correction. **d.** Basal OCR (**d**) and maximal OCR (**e**) showing significant decreases in the Δ*ctr2* and Δ*ctr2*:Ctr2-HA parasite lines. Bars are at mean ± SD of three independent experiments. *p* value from one-way ANOVA with Tukey’s correction. **f**. ECAR showing no significant changes in any parasite lines, bars are at mean ± SD of three independent experiments. **g.** Metabolic map demonstrating a small shift in parasite metabolism. Points at mean ± SD. **h**. Quantification of plaque number of parental, Δ*ctr2* and Δ*ctr2*:Ctr2-HA parasites after treatment with 5 µM CuSO_4_. Bars at mean ± SD. *p* values from one-way ANOVA with Tukey correction. **i.** Ctr2-HA adopts a more peripheral localisation in BCS-treated parasites. DNA in blue. Scale bar 5 μm. **j.** Quantification of Ctr2-HA localisation. Bars are mean ± SD of three independent experiments. *p* value from two-way ANOVA, Fisher LSD test. Total number of vacuoles (**k**) and cysts (**l**) at 48 h in pH 8.1 for indicated strains. Bars at mean ± SD. *p* values from one-way ANOVA with Tukey correction. **m**. % of vacuoles DBA positive after 24 h infection in neurons. Bars at mean ± SD. *p* values from one-way ANOVA with Tukey correction.

### Supplementary tables

**Table S1:** Summary of RNAseq data.

**Table S2:** Table of normalised data from quantitative metabolomic data, including all detected metabolites.

**Table S3**: Summary of quantitative metabolomics data.

**Table S4:** Table of raw data from stable isotope labelling, including all detected isotopomers, after ^13^C-glucose and ^13^C-glutamine labelling.

**Table S5**: Table of normalised stable isotope labelling data.

## METHODS

### Alignments

Protein sequences were identified using BLAST and VEuPathDB^92^ and full length alignment was performed using muscle5^93^. A maximum likelihood-based phylogenetic tree was generated using the *iqtree* default parameters for ModelFinder + tree reconstruction + ultrafast bootstrap (1000 replicates). The unrooted phylogenetic tree was visualized using ITOL^94^.

### *Toxoplasma gondii* and host cell maintenance

*Toxoplasma gondii* tachyzoites were grown in human foreskin fibroblast (HFFS) cultured in Dulbecco’s modified Eagle’s medium (DMEM), supplemented with 3% or 10% (D3, D10) heat-inactivated foetal bovine serum (FBS), 2 mM L-glutamine and 50 U/mL Penicillin/Streptomycin and maintained at 37°C with 5% CO_2_. Bradyzoites were maintained in Roswell Park Memorial Institute (RPMI) media, supplemented with 1% FBS and 50 U/mL Penicillin/Streptomycin.

### Copper depletion using BCS

To deplete available copper, HFF cells were split 1:4 in the presence of 100 µM Bathocuproinedisulfonic acid (BCS) for two passages. *T. gondii* parasites were then grown in BCS-treated host cells for one passage (48h) before the experiment, which was carried out in the presence of BCS.

### qRT-PCR

Parasites were treated as described, mechanically released and filtered through a 5 μm pore filter to remove host-cell debris and pelleted by centrifugation (2000 x g for 10 min), supernatant was removed, and parasite pellet was washed once in sterile PBS before spinning down in a microcentrifuge at 7000 x rpm for 10 min. Supernatant was removed and pellets were frozen at -20°C overnight. RNA was then extracted from the pellets using the RNeasy Mini Kit (Qiagen, 74104). 1 µg RNA was then treated with DNAse I 1 U/µL (ThermoFisher Scientific, 18068015) at room temperature for 15 mins before inactivating the DNAse with 25 mM EDTA followed by DNase denaturation at 65°C for 10 min. Using 1 µg of DNase I treated RNA, cDNA synthesis was then performed using the High-Capacity cDNA Reverse Transcription Kit (Applied Biosystems, A48571) according to manufacturer’s instructions. qRT-PCR was carried out on the Applied Biosystems 7500 Real Time PCR machine using PowerSYBR green PCR master mix (ThermoFisher Scientific 4367659) and 2 ng of 1:10 diluted cDNAs per reaction. Primers sequences are indicated in **Table 1**, and actin (P9 and P10) included as a housekeeping control. Results are from three independent biological experiments, each performed in technical triplicate. Relative fold changes were calculated using the Pfaffl method ^95^. GraphPad Prism 10 was used to perform statistical analysis.

**Table 1:**
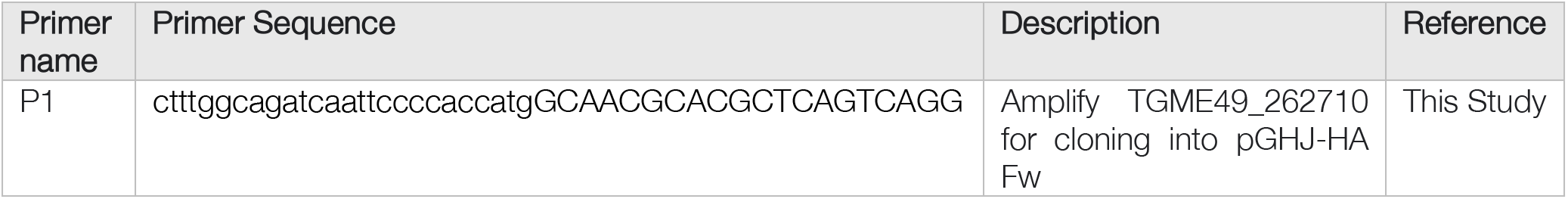
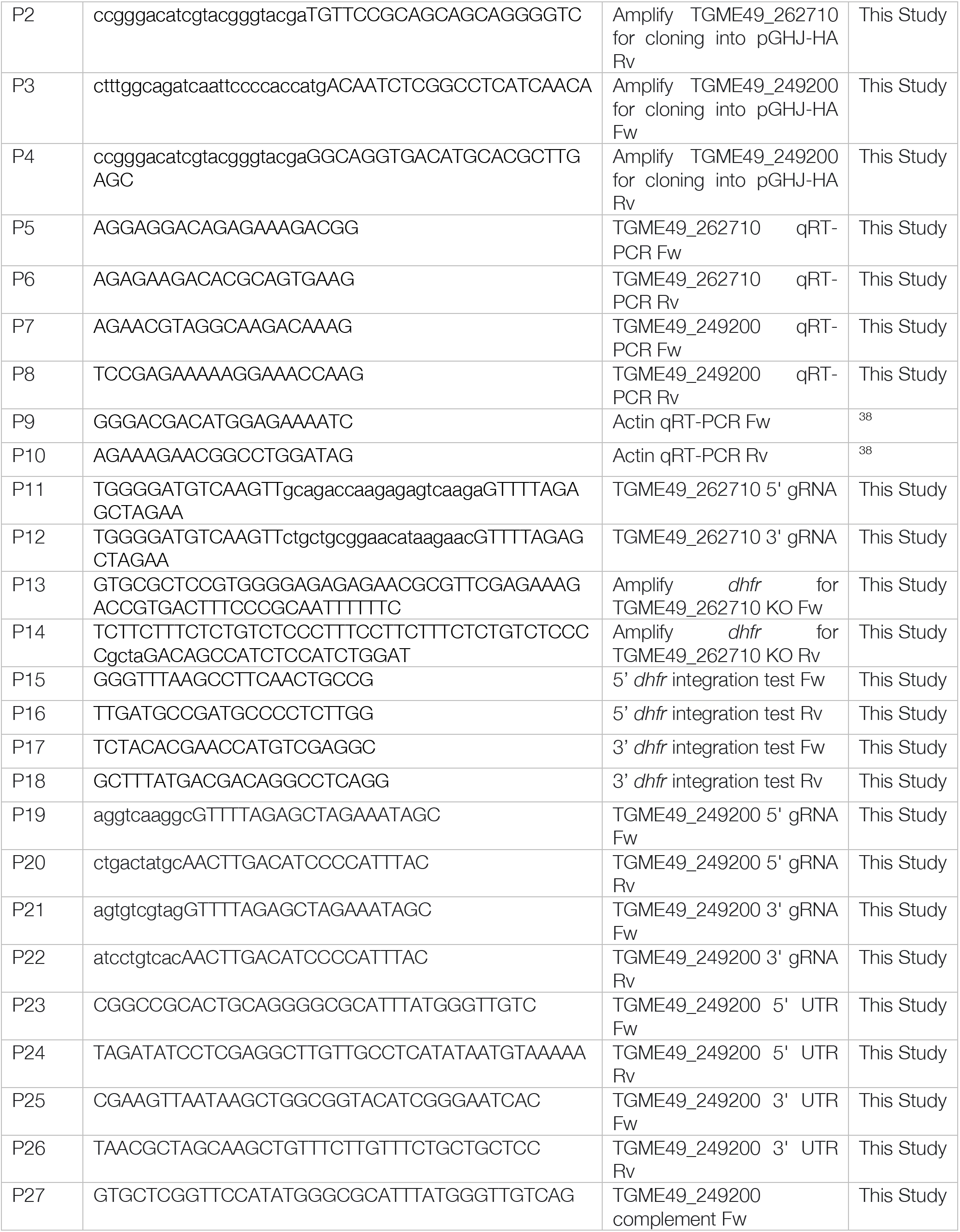
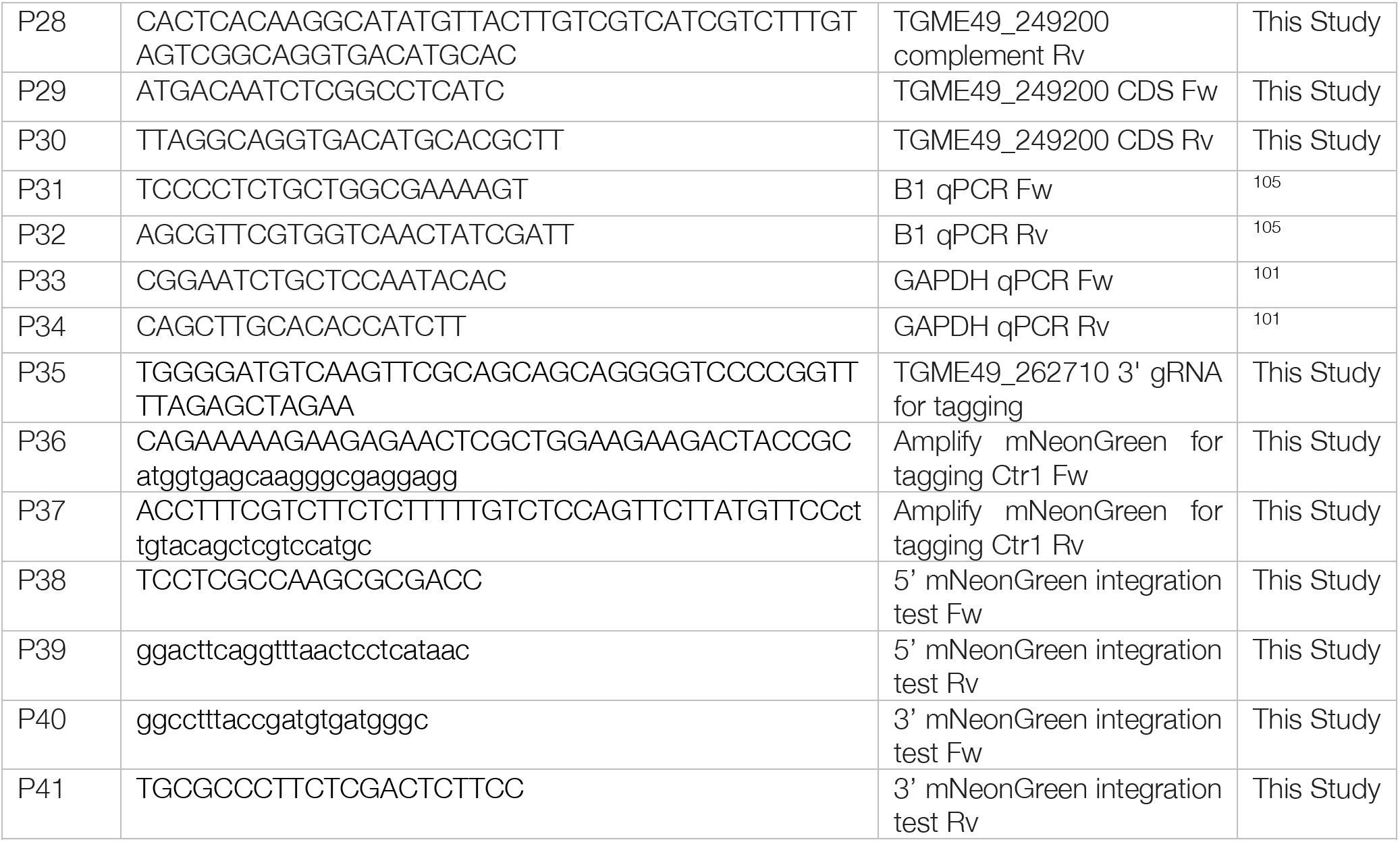
List of primer sequences used in this study.

### *Xenopus* oocytes ectopic expression

#### Ethics statement

*Xenopus laevis* frog maintenance and oocytes preparation was approved by the Australian National University Animal Experimentation Ethics Committee (Protocols A2014/20 and A2020/48). Before surgeries to extract oocytes, frogs were anesthetized by submersion in a 0.2% (w/v) tricaine methanesulfonate (MS-222) solution made in tap water and neutralized with Na_2_HCO_3_ for 15–40 mins until frogs exhibited no reaction upon being turned upside down.

#### Xenopus laevis oocytes preparation and Ctr1-HA expression

The open reading frame of *ctr1* (P1 and P2) or *ctr2* (P3 and P4) was amplified from RHΔ*Ku80* cDNA using the indicated primers. The resultant products were cloned by Gibson assembly into the vector pGHJ-HA previously digested with *Xma*I and *Avr*II enzymes^96^. The plasmids were linearized using *Not*I, and complementary RNA (cRNA) encoding HA-tagged *ctr1* or *ctr2* was prepared for injection into oocytes as previously described^97^. *Xenopus laevis* oocytes were surgically removed and prepared for cRNA injection as described^98^. 20 ng of *ctr1* or *ctr2* cRNA was micro-injected into stage 5 or 6 oocytes using a Micro4^TM^ micro-syringe pump controller and A203XVY nanoliter injector (World Precision Instruments).

#### Oocytes surface biotinylation and whole membrane preparation

*Xenopus laevis* oocytes surface biotinylation was performed as described previously^9,98^. Briefly, for surface biotinylation, 5 oocytes were selected 5 days post cRNA injection, washed three times with ice-cold PBS (pH 8.0), incubated for 10 min at room temperature in 0.5 mg/ml of EZ-Link^TM^ Sulfo-NHS-LC-Biotin (Thermo Fisher Scientific), and then washed four times with ice-cold PBS. Oocytes were subsequently solubilized in oocyte lysis buffer (20 mM Tris-HCl pH 7.6, 150 mM NaCl, 1% v/v Triton X-100) for 2 hr on ice. Samples were centrifuged for 15 mins at 16,000 *g*, and the supernatant was mixed with 50 μl of streptavidin-coated agarose beads (Thermo Fisher Scientific). The mixture was incubated at 4°C on slow rotation overnight. Beads were washed four times with oocyte lysis buffer before elution in 20 µL of SDS-PAGE sample buffer with 4% b-mercaptoethanol. Protein samples from surface biotinylation were separated by SDS-PAGE. Proteins were then wet transferred to nitrocellulose membrane (0.2 µM, Cytiva, 10600004) in Towbin buffer (0.025 M Tris, 0.192 M glycine, 10 % methanol) for 60 mins at 250 mA and blocked for at least 1 h at room temperature in blocking buffer (5 % milk in 0.1 % Tween/PBS). Blots were then stained with primary antibodies overnight at 4°C (rat anti-HA (1:200–1:400 dilution; clone 3F10 Sigma-Aldrich, cat. number 11867423001), followed by secondary horseradish peroxidase-conjugated goat anti-rat IgG (1:5,000 or 1:10,000 dilution; Abcam, cat. number ab97057) antibodies at room temperature for 1 h. Antibody-labeled membranes were incubated in enhanced chemiluminescence solution (0.04% w/v luminol, 0.007% w/v coumaric acid, 0.01% H_2_O_2_, 100mM Tris pH 9.35) and imaged using a ChemiDoc MP imaging system (Bio-Rad).

#### Oocyte copper uptake by ICP-MS

The uptake of copper into oocytes was measured 4 days post-cRNA injection. Batches of ten non-injected oocytes or oocytes expressing Ctr1-HA were washed four times with ND96 buffer (96 mM NaCl, 2 mM KCl, 1 mM MgCl2, 1.8 mM CaCl2, 10 mM MES; pH 5.5) at RT and then incubated in ND96 with or without 25 µM CuSO_4_ supplemented with 2 mM ascorbic acid for 30 min or 1 h. After incubation, oocytes were washed four times in 4 mL of ice-cold ND96, with the first wash using ND96 supplemented with 1mM EDTA. The oocytes were freeze dried, then lysed in 125 µL of ultrapure water; once fully lysed, 125 µL of 4% nitric acid was added into the samples. The samples were centrifuged at > 15,000 *× g* for 10 min and the supernatant was collected for analysis by ICP-MS. Each condition was performed in quintuplicate. Elemental analysis was performed using the Thermo Fisher iCAP RQ ICP-MS equipped with an autosampler, in the high sensitivity kinetic energy discrimination (KEDS) mode. The 65Cu isotope was measured to determine Cu content in oocyte samples. Calibration standards were prepared from a multi-element standard solution containing Cu (Standard No. 1 for ICP-OES; Agilent). A quality control reference consisting of the calibration standard solution at an element concentration of 10 ppb was measured before the beginning of the sample run, after every 10–12 samples, and at the end of the sample run to correct for variation between and within ICP-MS analysis runs. Sample concentrations were calculated using external calibration methods within the instrument software. Further data processing was performed in Microsoft Excel.

#### Oocyte copper uptake by PhenGreen quenching

PhenGreen quenching was performed as previously described^40,99^. Non-injected or 4 days post cRNA injection *Xenopus laevis* oocytes were washed 4 times with ND96 buffer at RT and then incubated with or without 20 µM CuSO_4_ (with 2 mM ascorbic acid in ND96 pH 5.5) for 1 h. Oocytes were then injected with 30 nL of 5 mM PhenGreen FL (Invitrogen, P6763), and incubated for 30 mins, in ND96 media. Oocytes were then washed with PBS, and pool of five oocytes were lysed in 100 µL of distilled water and pelleted for 10 min. PhenGreen fluorescence signal was measured in 100 µL of supernatant (Ex= 490 nm, Em= 520 nm) using a Tecan infinite M1000Pro plate reader in a 4x4 multiple reads per well matrix. Values were normalized to PhenGreen FL injected oocytes that were not incubated with copper.

### *Toxoplasma gondii* strain construction

*Δctr1:* 5’ and 3’ sgRNA sequences (P11 and P12, **Table 1**) for *ctr1* (TGME49_262710) were cloned into the pU6-cas9 plasmid, previously linearised using BsaI, by Gibson assembly. DHFR sequence was amplified by PCR using primers P13 and P14 containing tails with homologous sequences for the 5’ and 3’ UTR of *ctr1* (**Table 1**). The plasmids containing the 3’ and 5’ sgRNA sequences were transfected together with PCR amplified DHFR sequence into RHtdTomato parasites^38^. Parasites were selected using pyrimethamine and cloned by limiting dilution. Clones were confirmed by PCR using primer P15 and P16 for 5’ integration and primers P17 and P18 for 3’ integration.

*Δctr2:* 5’ (P19 and P20) and 3’ (P21 and P22) sgRNA sequences (**Table 1**) for *ctr2* (TGME49_249200) were cloned into the sgUPRT plasmid as previously described ^100^. A cassette encoding the *hypoxanthine phosphoribosyltransferase* (HXGPRT) gene was cloned into the pTKO vector^101^. This was flanked between homology regions targeting the upstream (using primers P23 and P24) and downstream (P25 and P26) untranslated regions, adjacent to the 3’ and 5’ Cas9 cut sites in *ctr2* gene. This plasmid, and the sgRNA sequences were transfected together into PruΔ*hpt* parasites. Correct integration of the resistance cassette was selected for using 25 mg/ml mycophenolic acid (MPA) and 50 mg/ml xanthine and parasites cloned by limiting dilution. Clones were confirmed by PCR using primers P29 and P30 for the absence of *ctr2* gene.

*Δctr2:*Ctr2-HA: TGME49_249200 was amplified from genomic DNA using primers 27 and 28, including a HA sequence, in frame at the 3’ end. The amplified sequence was cloned into a Bleo vector, transfected into the Δ*ctr2* parasite line. Selection for bleomycin resistance was performed as previously described^102^, followed by cloning and complementation was confirmed by PCR using primers 27 and 28.

Ctr1-mNGreen: sgRNA sequence (P35, **Table 1**) for *ctr1* (TGME49_262710) was cloned into the pU6-cas9-YFP plasmid^103^, previously linearised using BsaI, by Gibson assembly. mNeonGreen sequence was amplified by PCR using primers P36 and P37 containing tails with homologous sequences for *ctr1* (**Table 1**). The plasmid containing the sgRNA sequence was transfected together with PCR amplified mNeonGreen sequence into RHΔKu80 parasites^38^. YFP positive parasites were sorted in a 96 well plate and checked by PCR for mNeonGreen integration. Clones were confirmed by PCR using primer P38 and P39 for 5’ integration and primers P40 and P41 for 3’ integration.

### *T. gondii* transfection

Parasite transfection was performed as previously described^104^. Parental parasites (RHΔKu80:tdTomato or PruΔhxgprt) were mechanically lysed, filtered through a 5 μm pore filter to remove host-cell debris and pelleted by centrifugation (1,500 rpm for 7 min). Pelleted parasites were then resuspended in 400 μL cytomix electroporation buffer (10 mM K_2_HPO4/KH_2_PO_4_, 120 mM KCl, 150 μM CaCl2, 5 mM MgCl2, 25 mM HEPES and 2 mM EGTA), supplemented with 2 mM ATP and 5 mM GSH. Electroporation was performed using an ECM 830 Square Wave electroporator (BTX) in 4 mm cuvettes. Following electroporation, parasites were placed onto confluent HFFs and selected as described above.

### Plaque assay

Confluent HFFs in 6 well plates were infected with 250-500 mechanically lysed tachyzoites of the indicated strain in D3 or D3 with supplements (5 or 20 μM CuSO_4_, 250 µM uracil, 200 μM ferric ammonium chloride (FAC), 25 μM ZnSO_4_, concentrations based on^9^) as indicated and incubated for 7-10 days. Media was then removed, washed with PBS, and fixed in ice cold methanol for 10-20 min. Fixed monolayers were stained with crystal violet (12.5 g crystal violet in 125 ml ethanol, diluted in 500 ml 1% ammonium oxalate) at room temperature and subsequently washed with deionized water.

### Extracellular survival assay

Extracellular survival was quantified as previously described^38^. Briefly, parasites were mechanically released, counted and diluted in D3 media. 500 parasites per well of a 6 well plate were added immediately (for time 0) and the remaining incubated in D3 at 37 °C until the indicated timepoints. Cells were fixed and stained as above after 8 days of incubation, and plaque numbers counted.

### Immunofluorescence microscopy

Confluent HFFs grown on 13 mm glass coverslips were infected with 10,000 - 250,000 mechanically lysed tachyzoites. At the indicated time, media was removed and cells fixed with 4% paraformaldehyde for 20 min at room temperature. Cells were permeabilized and blocked for 1 hr at room temperature with 0.1% Triton X-100 in phosphate buffered saline (PBS) supplemented with 3% bovine serum albumin (BSA). Cells were incubated with primary antibody (**Table 2**) in blocking buffer (0.1% Triton X-100, 1% BSA in PBS) for 1 hr at room temperature or overnight at 4°C. Cells were then washed three times with PBS and incubated with secondary antibodies (**Table 2**) in blocking buffer. The coverslips were then either incubated with 1:1000 4′,6-diamidino-2-phenylindole (DAPI) in PBS followed by 3 washes and mounted using Prolong Diamond (Life Technologies, P36970) or mounted directly using Fluoromount with DAPI (Southern Biotech, 0100-20) and allowed to set overnight at room temperature. Parasites were imaged using a Leica DiM8 (Leica Microsystems) microscope and processed using LasX (Leica Microsystems). Intensity was analysed using automated ImageJ macros (Fiji).

**Table 2.** List of antibodies used for IFA.

| **Antibody** | **Concentration** | **Reference** |
|---|---|---|
| Rabbit anti-*T. gondii* polyclonal | 1:10,000 | Invitrogen (PA17252) |
| Dolichos biflorus agglutinin (DBA) | 1:500 | Vector Laboratories (1031) |
| Rabbit anti-HA | 1:500 | CST (C29F4) |
| Mouse anti-HA | 1:500 | CST (6E2) |
| Rat anti-HA | 1:500 | Sigma (11867423001) |
| Rabbit anti-Sag2x | 1:1,000 | <sup>106</sup> |
| Rabbit anti-SRS9 | 1:500 | <sup>107</sup> |
| Mouse anti-Sag1 | 1:10,000 | Boothroyd et al., 1997 |
| Mouse anti-F1BATPase | 1:3,000 | <sup>108</sup> |
| Rat anti-DRPB | 1:1,000 | <sup>109</sup> |
| Mouse anti-SERCA | 1:5,000 | <sup>110</sup> |
| Guinea pig anti-CDPK1 | 1:10,000 | <sup>61</sup> |
| Rabbit anti-TOM40 | 1:1,000 | <sup>59</sup> |
| Mouse anti-mNgreen | 1:1,000 | Proteintech Cat # 32f6 |
| Goat anti-Mouse HRP | 1:10,000 | Invitrogen Cat # 31430 |
| Alexa Fluor goat anti-rabbit 488 | 1:500 | Invitrogen Lot #2891813 |
| Alexa Fluor goat anti-mouse 488 | 1:500 | Invitrogen Lot #2486523 |
| Alexa Fluor goat anti-guinea pig 488 | 1:500 | Invitrogen A-11073 |
| Alexa Fluor goat anti-rat 488 | 1:500 | Invitrogen A-11006 |
| Alexa Fluor goat anti-rat 568 | 1:500 | Invitrogen Lot #1853640 |
| Alexa Fluor goat anti-mouse 568 | 1:500 | Invitrogen Lot #2447869 |
| Alexa Fluor goat anti-rabbit 568 | 1:500 | Invitrogen Lot #2856117 |
| Alexa Fluor goat anti-guinea pig 568 | 1:500 | Invitrogen A-11075 |

### Toxicity Assays

Toxicity assays of fluorescent parasites were performed as previously described^38^. Briefly, host cells were infected with 1,000 either RHtdTomato or RHtdTom Δ*ctr1* tachyzoites per well and allowed to invade for 2-4 hours at 37°C. The media was then removed and supplemented with CuSO_4_ (at indicated concentrations) and incubated for 4-6 days. Total parasite fluorescence was recorded using a PHERAstar FS microplate reader (BMG LabTech). All experiments were performed in triplicate with at least three independent biological replicates. Uninfected wells served as blanks and fluorescence values were normalised to infected, untreated wells. Dose-response curves and EC_50_s were calculated using GraphPad Prism 10.

### Growth Assay

To assess parasites replication, parasites were allowed to infect HFF cells grown on coverslips for 24 or 48 h prior to fixation as above. Where required, parasites were stained using primary rabbit anti-*T. gondii* or anti-CDPK1 (**Table 2**) and appropriate secondary antibodies. At least 100 vacuoles were counted manually and scored by the number of parasites.

### ICP-MS

Total parasite associated copper, iron and zinc were quantified using ICP-MS. Confluent HFFs cells in T150 flasks were infected with corresponding parasites and let them grow for 3 days. Parasites were harvested by scraping, syringing and filtering through either 0.3 µM filters or PD-10 columns (Cytiva, 17085101), and quantified using a haemocytometer. Parasites were then washed three times with PBS supplemented with 6% Chelex 100 sodium form (Merck, C7901), then once with 1 mM EDTA before being digested in 65% nitric acid (Merck, 225711) at 85°C for 5 h. Digested samples were then diluted 1:10 in 2% nitric acid with 0.025% Triton X-100. At least three biological replicates were collected, with three technical replicates per biological sample. Samples were quantified at the Scottish Universities Environmental Research Centre (SUERC) and processed by Dr Valerie Olive. Briefly, samples were run on an Agilent 7500ce ICP-MS, fitted with a double pass spray chamber and a self-aspire nebulizer tuned for 0.1 ml/min. A 3 ppb Indium (115In) was introduced as internal standard to correct for the sensitivity drop caused by the sample matrix. Cu and Zn were taken on the masses 63 and 66 respectively and the results are an average of 10 repeats. The calibration equation was calculated from 3 points at 5, 10 and 20 ppb using Cu and Zn mono-element solutions Cu and Zn specpure ®. Iron was taken at the mass 56 using a collision cell with a flow of Hydrogen at 4 ml/min to suppress the interference of the 40Ar16O. The calibration equation was calculated using a 200 ppb and 300 ppb solution of mono-element Fe specpure ®, results are an average of 20 repeats.

### X-ray fluorescence microscopy (XFM)

#### Sample preparation

Cell preparation was performed as described^111^. HFF primary cells (HFF-1 (ATCC® SCRC-1041TM)) were cultured directly on Si_3_N_4_ membranes (Silson, UK). The HFF cells were infected with parental or mutant *T. gondii* after 24h. Vitrification was performed 24h post-infection with EasyGrid^112^. Samples were rinsed in a 150 μM acetate solution to remove buffer salts and then placed in the sample holder of EasyGrid and pressure wave generation used to remove excess liquid. Cryocooling performed by jetting liquid propane on both sides of the membrane.

#### Data collection and analysis

Samples were transferred to the cryo-stage of the ESRF beamline ID16A using EM-VCM and EM-VCT (Leica systems) to be imaged on the cryostage at 120K under high-vacuum. Samples were scanned using “on-the-fly” acquisition modes in which the sample is horizontally translated at constant speed^113,114^ with a focussed beam of 20 x 20 nm. The fluorescence signal emitted from each sample pixel was recorded by 2 custom multi-element Silicon Drift Detectors (SDD) placed on both sides of the sample at 90 degrees from the incident X-ray beam. A multi-element SDD (Hitachi Ltd.) and HARRY coupled to ARDESIA-16 spectrometers based on a monolithic SDD array^115^ were used. Nanometer scale 2D sample thickness maps were also measured for each prepared Si_3_N_4_ membrane. Fluorescence scans were then performed at a spatial resolution of 50 nm (step size) using a dwell time of 50 ms at high dose. The resulting XRF spectra were fitted pixel by pixel using an in-house Python script based on PyMca libraries developed at the ESRF^116^ to correct for detector deadtime and to normalise the incident X-ray beam variation. The resulting elemental area mass density maps were analysed with ImageJ.

### RNAseq

Parental (RHtdTomato), Δ*ctr1*, or Δ*ctr1* grown in the presence of 600 µM copper for 24 h were mechanically released, collected by centrifugation at 1500×*g* for 10min and pellets stored at −80°C until required. RNA was extracted from the pellets using the RNAeasy kit (Qiagen) according to the manufacturer’s instructions. RNA libraries were then prepared using Illumina Stranded mRNA library preparation method and sequenced at 2×75bp to an average of more than 5 million reads per sample. Raw sequencing data (FASTQ format) was processed using the Galaxy public server hosted by EuPathDB (https://veupathdb.globusgenomics.org/). FastQC and Trimmomatic were used for quality control and to remove low quality reads (where Q < 20 across 4bp sliding windows) and adaptor sequences^117^. The filtered reads were aligned to the *T. gondii* ME49 genome using HITSAT2^118^. These sequence alignments were used to identify reads uniquely mapped to annotated genes using HTSeq-count. Differential expression analysis was performed in R using DESeq2^119^ and pheatmap. All processed data are available in **Table S1** and raw FASTA files are available at http://www.ncbi.nlm.nih.gov/bioproject using bioproject ID: PRJNA1387371.

### MitoTracker Imaging

Parasites treated as above were mechanically released, washed once with PBS and incubated in the presence and 50 nM MitoTracker Green (Invitrogen, M46750) for 10 min at 37 °C on poly-L-lysine treated glass bottom dishes. Parasites were imaged live immediately using Leica DMi8 (Leica Microsystems) microscope and processed using LasX (Leica Microsystems) and FIJI.

### Seahorse Assay / Extracellular flux analysis

*Toxoplasma* Seahorse assay was performed as previously described^57^. Briefly, parasites were incubated in the absence or presence of 20 µM of CuSO_4_ for 24 h, extracellular parasites removed by washing 1X PBS, and intracellular parasites were mechanically released, filtered and harvested by spinning at 2000 x g for 10 min. Parasites were then resuspended in Seahorse XF media (Agilent) supplemented with 5 mM glucose and 1 mM L-glutamine (Sigma-Aldrich, G7513) at 37°C. 1.5 x 10^6^ parasites in a final volume of 175 µL were added to each well of a poly-L-lysine-treated seahorse XF cell culture miniplate (Agilent). Parasites were then centrifuged at 300 x g for 5 mins to ensure sedimentation before the basal oxygen consumption rate (OCR) and the extracellular acidification rate (ECAR) was quantified using a Seahorse XF HS Mini Analyser (Agilent Technologies). The OCR following the addition of 1 µM atovaquone (Scientific, 17575545) was considered as the non-mitochondrial oxygen consumption rate and subtracted as background. The basal and maximal oxygen consumption rates of the parasites were calculated by subtracting the background from the raw OCR values before and after the addition of 1 µM Carbonyl cyanide 4-(trifluoromethoxy) phenylhydrazone (FCCP) (Merck, C2920) respectively. Basal ECAR values were taken as raw values. This experiment was performed in three independent biological replicates, each in triplicate.

### Clear-native PAGE and Complex IV activity assay

Clear-native PAGE and complex IV activity stain was performed as described previously^60^. Briefly, whole parasite samples were suspended in solubilisation buffer (50 mM NaCl, 50 mM imidazole, 2 mM 6-aminohexanoic acid, 1 mM EDTA–HCl pH 7.0, 2% (w/v) n-dodecyl-maltoside, 1% digitonin) and incubated on ice for 10 min followed by centrifugation at 16,000 × g at 4°C for 15 mins. The supernatant containing solubilised membrane proteins was combined with glycerol and ponceau S to a final concentration of 6.25% and 0.125% respectively. Samples (equivalent of 1.5 x 10^7^ parasites per lane) were separated on a NativePAGE 4–16% Bis-Tris gel. NativeMark was used as a molecular weight marker. In case of protein detection by Western blot, proteins were wet transferred to a PVDF membrane in Towbin buffer (0.025 M Tris, 0.192 M glycine, 10 % methanol) for 60 mins at 250 mA and blocked for at least 1 h at room temperature in blocking buffer (5 % milk in 0.1 % Tween/PBS). Blots were then stained with primary antibodies overnight at 4°C (mouse anti-mNgreen), followed by Anti-mouse IgG secondary horseradish peroxidase-conjugated secondary antibody at room temperature for 1 h. Ctr1-mNgreen was detected using Pierce ECL Western Blotting Substrate (ThermoFisher Scientific) and imaged using the Invitrogen iBright 1500 Imaging System. For complex IV oxidation activity, the gel was incubated in a 50 mM KH_2_PO_4_, pH 7.2, 1 mg/ml cytochrome c (Sigma-Aldrich, C2506), 0.1% (w/v), 3,3’-diaminobenzidine tetrahydrochloride solution (Sigma-Aldrich, D5637) for 24 h.

### Western blotting

Parasites were mechanically released, filtered, counted in haemocytometer chamber and collected by centrifugation at 1500 x g for 10 min, and lysed with RIPA lysis buffer (150 mM sodium chloride, 1 %Triton X-100, 0.5 % sodium deoxycholate, 0.1 % sodium dodecyl sulphate (SDS) and 50 mM Tris, pH 8.0) in a ratio of 10^6^ parasites/µL for 30 mins on ice. Samples were then resuspended with 5X Laemmli buffer (10% SDS, 50% glycerol, 300 mM Tris-HCL pH 6.8 and 0.05% bromophenol blue), boiled at 95°C for 5 min (unless stated otherwise) and separated on a 10% SDS-PAGE gel. PageRuler Prestained Protein Ladder (ThermoFisher Scientific, 26619) was used as a molecular weight marker. Proteins were then wet transferred to nitrocellulose membrane (0.2 µM, Cytiva, 10600004) in Towbin buffer (0.025 M Tris, 0.192 M glycine, 10 % methanol) for 60 mins at 250 mA and blocked for at least 1 h at room temperature in blocking buffer (5 % milk in 0.1 % Tween/PBS). Blots were then stained with primary antibodies overnight at 4°C (rabbit anti-TOM40^59^, a kind gift from Giel van Dooren), followed by Anti-Rabbit IgG secondary horseradish peroxidase-conjugated secondary antibody (Promega, W401B) at room temperature for 1 h. Specific signals were detected using Pierce ECL Western Blotting Substrate (ThermoFisher Scientific) and imaged using the Invitrogen iBright 1500 Imaging System.

### ATP quantification

ATP was quantified as previously described^24^. Briefly, infected cells were washed twice with PBS, and FluoroBrite (Thermo Fisher Scientific, A1896702) supplemented with 1% FBS and HALT protease inhibitors (Thermo Fisher Scientific, 78430) was added. Cells were scraped and mechanically lysed and parasites were filtered through a 5 μm pore filter to remove host-cell debris and pelleted by centrifugation (1000 x g for 10 min). Parasites were resuspended in glucose- and glutamine-free Seahorse XF media (Agilent cat. N° 103575) and counted using a haemocytometer chamber. Parasites were added to a 96 well plate (1.5 x 10^6^ parasites/well in a final volume of 50 µL) and 50 µl of the corresponding compounds diluted in glucose- and glutamine-free Seahorse XF media were added accordingly to each well (5 mM 2-DG (Sigma-Aldrich, D6134), 25 mM glucose, 2 mM L-glutamine (Sigma-Aldrich, G7513) or 20 µM oligomycin (Sigma-Aldrich, 495455)) and incubated for 1 h. The reaction was stopped by flash freezing samples in liquid nitrogen. After thawing, 100 µL of CellTiter-Glo reagent (Promega, G7572) was added directly to the wells and incubated for 1 h. Relative ATP levels were determined by normalizing luminescence to untreated parental parasites, data analysis performed using GraphPad Prism 10.

### Glucose uptake assay

Glucose uptake was assessed by flow cytometry^58^ using the fluorescent glucose analogue 2-NBDG (Invitrogen, N13195). Parasites were mechanically released and filtered to remove host material. After washing, parasites were resuspended in DMEM supplemented with 2.5 µM L-glutamine (Sigma-Aldrich, G7513) or 2.5 µM L-glutamine and 60 mM glucose (for unlabelled competition). Parasites were then left unstained or were incubated with 1 mM 2-NBDG for 30 mins at 37°C. Fluorescence was analysed on a BD FACSCelesta Flow Cytometer and data was acquired using FACSDiva software (BD Biosciences). Parasites were gated on forward and side scatter and red fluorescence (only for TdTomato parasites). All data were analysed using FlowJo v10 (BD Biosciences) and the geometric mean of the BB515 channel was used to quantify 2-NBDG uptake. Each data point was normalised to the sum of its replicate to both show variation among replicate while also controlling for variation in total fluorescence between them ^120^.

### Untargeted Metabolomics sample preparation

For metabolite extraction, 4 x 15 cm dishes for each condition (control, Δ*ctr1* or Δ *ctr1* incubated with 20 µM CuSO_4_ for 24 h) were pooled to generate each replicate. After 48 hours of infection, cells were washed with 20 ml of ice-cold PBS twice to remove extracellular parasites and old media, before being scraped and passed through a 25-gauge needle to mechanically release parasites. Parasites were filtered from host material through a 3 µm filter before centrifugation at 1°C for 25 mins at 3000 x g. Pellets were resuspended and washed in 1 ml ice-cold PBS before being counted using a haemocytometer. Parasites were resuspended in 200 µl of a chloroform:methanol:water mixture (1:3:1 ratio). Samples were shaken for 1 hour at 4°C and centrifuged at 20,000 x g for 5 mins at 4°C. Metabolites in the supernatant were then collected.

### Metabolomics data analysis

Metabolites were quantified by LC-MS as previously described^121^. Samples were separated by high-performance liquid chromatography on a Dionex UltiMate 3000 RSLC system (Thermo) using a ZIC-pHILIC column (Merck). Mass spectrometry was performed using an Orbitrap Q Exactive (Thermo) with analysis performed in both positive and negative ionization modes. Internal standard mixtures, consisting of metabolites from a range of metabolic pathways were run to aide in metabolic annotation. Fragmentation of pooled control samples to obtain MS2 spectra using data dependent acquisition (DDA) was also performed. Features were annotated using the mzMatch package in R along with the IDEOM software^122,123^. A root squared deviation (RSD) filter was applied and features with greater than 40% in QC samples were removed, along with those with retention times < 3 min. Metabolites were annotated according to the Metabolite Standards Initiative^124^ with level 2 (mass only) and level 1 annotations included in downstream pathway analysis. MSI level 1 annotations correspond to metabolites present in the mixture of 240 standards run alongside experimental samples. Peaks were also subjected to manual filtering to remove poor-quality peaks. Features identified in only one condition have missing values imputed with 1/5 the minimum value detected. Median normalization, log10 transformation, and mean centering were performed on peak intensities from identified metabolites prior to hierarchical clustering and differential abundance analysis within MetaboanalystR 4.0^125^.

Stable isotope labelling analysis was performed in RStudio as described previously^58^. Briefly, AccuCor corrected isotopologue intensities were normalized to the proportion of total metabolite intensity^126^. To test for changes in isotopologue distributions, data were first centred log-ratio (CLR) transformed to mitigate for dependencies in the normalized data. Labelling extent (LE) (1-M0 fraction) was calculated for each metabolite^119^ and used in principal components analysis. For pathway heatmaps, KEGG compound IDs were used to identify metabolic pathways associated with each metabolite. The average LE for metabolites across conditions was determined, and heatmaps were then generated for the listed pathways. In the rescue analysis, LE values were logit-transformed to ensure their suitability for linear modelling before differences in labelling abundance were tested between parental and Δ*ctr1* using the limma linear model^127^. *p* values were then corrected for multiple testing using the Benjamini-Hochberg false discovery rate (FDR), with metabolites with an FDR ≤ 0.01 being retained for rescue analysis. For this subset of metabolites, we calculated a signed rescue score which represents the fraction of the change between parental and Δ*ctr1* that is reversed by the addition of copper in our Δ*ctr1*+Cu condition.

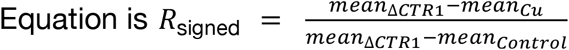

In this analysis, a value of 1 indicates full rescue, 0 indicates no rescue and values greater than 1 or less than 0 represent overcompensated or exacerbated phenotypes respectively. Rank ordered rescue scores were expressed as a percentage before being plotted using GraphPad prism. Raw metabolomics mass spectrometry data are available at the MetaboLights (https://www.ebi.ac.uk/metabolights/) data repository under project accession MTBLS13492.

### Cyst assay

Cyst assays were performed and analysed as previously described^64^. Confluent HFF monolayers were grown in 96 well plates (Perkin Elmer; 6055302) and inoculated with mechanically-lysed tachyzoites of indicated strain, followed by spinfection. For the alkaline stress model of bradyzoite culture, HFFs were infected with parasites for 4-6 hours under non-cyst inducing conditions (pH 7.1 and 5% CO2). After 24 hours, the media was changed to alkaline media (50 mM HEPES [pH 8.1] in RPMI supplemented with 1% fetal bovine serum, penicillin, and streptomycin). After the indicated time point of infection, plates were fixed and stained using anti-*T. gondii* and biotinylated *Dolichos biflorus agglutinin* (DBA), a lectin which binds to the cyst wall (Vector Laboratories 1031, 1:500) followed by incubation with anti-rabbit Alexa Fluor 488 antibody and Streptavidin 647. Stained plates were imaged using Operetta High-Content Microscopy and analysed using Harmony Analysis software.

### Abnormal cyst quantification

To quantify the incidence of abnormal cyst formation across our parasite strains, we again performed cyst assays as described above. Plates were imaged and analysed using Harmony Analyzer software. A novel protocol was developed for abnormal cyst analysis, which built onto existing cyst analysis pipelines. After cysts were identified according to the original pipeline, they were then subjected to a symmetry, threshold, axial ratio, and roundness (STAR) analysis, which provides the software with extensive descriptions of cyst morphology and differentiation of phenotypes. Following STAR analysis, Harmony’s machine learning software was trained using 10 manually classified cysts of each phenotype to recognize normal and abnormal cysts. This pipeline would then quantify the number of abnormal cysts and normal cysts in each field of view, to generate a total number of each per strain. The number of abnormal cysts was divided by the total number of cysts and multiplied by 100 to generate a % abnormal cyst value for each strain.

### Inductively Coupled Plasma Optical Emission Spectroscopy (ICP-OES)

Confluent HFFs in T75 flasks were infected with tachyzoites from two fully lysed T25 flasks. At 1dpi, infected HFFs were scraped, mechanically lysed by passage through 25-27-gauge needles, filtered through a 5 μm pore filter to remove host-cell debris and pelleted by centrifugation (1,500 rpm for 7 min). Trace metal grade HNO_3_, at 70% (Macron 6623-46) was added to pellet samples followed by overnight incubation at 65°C. After incubation, the lysed pellets were diluted to 2.5% HNO_3_ by adding ultra-pure water (18.1 MΩ). Liquid samples were analysed for metal content using an iCAP PRO XDUO ICPOES with a wavelength of 324.754 nm for copper, 213.856 nm for zinc, 259.373 nm for manganese, 259.940 nm for iron, 279.55 nm for magnesium and 396.84 nm for calcium. ICP calibration standards (Inorganic Ventures) were prepared using IV-STOCK-27 and metal content was calculated using the Qtegra software.

### Reactivation assay

Confluent HFFs in 6 well plates were infected with 250 mechanically lysed tachyzoites of the WT, *Δctr2* line and *Δctr2:*Ctr2-HA for 6 h under non-cyst inducing conditions (pH 7.1, 37°C and 5% CO_2_). After the incubation, the media was replaced with alkaline media (pH 8.1) and were incubated under cyst inducing conditions (37°C and 0% CO_2_) for 6 days. The plates were switched back to non-cyst inducing conditions for another 14 days for generation of plaques. The plates were stained for plaques as described above.

### Ethics statement

All mouse studies and breeding were carried out in strict accordance with the Public Health Service Policy on Humane Care and Use of Laboratory Animals and approved by the University of Arizona’s Institutional Animal Care and Use Committee (#12-391). All mice were bred and housed in specific-pathogen-free University of Arizona Animal Care facilities with a 14 hr/10 hr light/dark cycle, with ambient temperature between 68° and 75° F, and 30–70% humidity.

### Primary murine neuron culturing

Mouse primary cortical neurons were harvested from E17 mouse embryos obtained from pregnant Cre reporter. Dissections of E17 cortical neurons were performed as described previously^128^. Primary neuronal cell cultures were generated by methods described previously with minor modifications^128^. Culture plates were prepared by coating with 0.001% poly-L-lysine solution (Millipore Sigma, P4707, diluted in water 1:10) for plastic surfaces and 100μg/ml poly-L-lysine hydrobromide (Sigma, P6282, dissolved in borate buffer, pH 8.4) for glass surfaces overnight. They were washed three times water and transferred to plating media (MEM, 0.6% D-glucose, 10% FBS). Four hours after plating, media was exchanged to neurobasal media (Neurobasal base media, 2% B27 supplement, 1% L-glutamine and 1% penicillin-streptomycin). On day *in vitro* (DIV) 4, neurons received a half volume media change of neurobasal media with 5 μM cytosine arabinoside (AraC, Millipore Sigma, C6645) to inhibit glial proliferation. One third media exchanges with neurobasal media occurred every 3–4 days thereafter. All experiments were performed on 10 DIV neurons.

### Neuron cyst assay

Cyst assays were performed and analysed as previously described^128^. Primary neurons were cultured in 96 well plates (Perkin Elmer; 6055302) and inoculated with mechanically lysed tachyzoites of indicated strain, followed by spinfection. After 1 dpi, plates were fixed and stained as described above. Plates were imaged using Operetta High-Content Microscopy and analysed using Harmony Analysis software.

### *In vivo* infection and tissue collection

*In vivo* infection and tissue preparation was performed as previously described^65,129^. Mice were intraperitoneally injected with 10,000 freshly syringe-lysed parasites of indicated strains in 200 μl of USP grade phosphate buffered saline (PBS). After 3 wpi, mice were euthanized by CO_2_ asphyxiation and perfused with 4°C PBS. Mouse brains were then harvested and were divided sagittally into left and right hemispheres. The left hemisphere was drop-fixed in 4 % paraformaldehyde (PFA) in PBS overnight at 4°C, rinsed in PBS, embedded with 30 % sucrose, and kept at 4°C prior to sectioning. After fixation brains were sectioned into 40 µm thick sagittal sections using a freezing sliding microtome (Microm HM 430). Sections were divided into 12 sequential tubes, so that each tube contained every 12th section. This ensured that each tube contained a relatively similar number of sections from equivalent regions of the brain. Sections were submerged in cryo-protective media (0.05 M sodium phosphate buffer containing 30% glycerol and 30% ethylene glycol) and stored at -20°C until staining. The right hemisphere of the brain was divided in half coronally, with the anterior half being flash-frozen and stored at -80°C until used for DNA extraction.

### Parasite burden by qPCR

To analyse the dissemination of parasites to the murine CNS in vivo, genomic DNA was extracted from the rostral half of flash-frozen brain hemispheres isolated from *T. gondii*-infected mice using the DNeasy Blood and Tissue kit (Qiagen, 69504). The *T. gondii* B1 gene was selected as the target of amplification given its high level of conservation and 35-fold repeat sequence within the parasite genome. Importantly, this gene is not expressed and is thus an accurate indicator of the number of parasite genomes present in the murine brain, rather than the number of mRNA transcripts, as is typically analysed by qRT-PCR. The B1 gene was amplified using primers 31 and 32^65^ (**Table 1**) in the presence of SYBR Green fluorescent dye, using the Eppendorf Mastercycler ep Realplex 2.2 system. Mouse GAPDH (primers 33 and 34 (**Table 1**)) was used as a housekeeping gene to normalize parasite DNA levels. The reaction conditions were as follows: 2 min at 50°C, 2 min at 90°C, followed by 40 cycles of 15 sec at 95°C, 15 sec at 58°C, and 1 min at 72°C, followed by a melting curve analysis.

### Cyst Counts *in vivo*

Cyst assays *in vivo* were performed as previously described^65^. Brain sections were washed in tris-buffered saline (TBS) and blocked in 3% Goat Serum in 0.3% TritonX-100/TBS for 1 hr. Sections were incubated overnight at room temperature with biotinylated DBA and rabbit anti-*T. gondii* polyclonal antibody (**Table 2**), washed, and incubated with the secondary antibodies streptavidin-Cy5 (Invitrogen, 1:500) and goat anti-rabbit 568 (1:500), DAPI (1:1000). 8 sections from each brain were mounted on glass slides using Fluoromount-G^TM^. The number of cysts (DBA and anti-*T. gondii* antibody positive) in each section were then quantified using a Revolve epifluorescent microscope (ECHO, San Diego). The sum of cysts from all 8 sections was combined to compare cyst burden across mice infected with different parasite strains.

## Data availability

All processed RNAseq data are available in **Table S1** and raw FASTA files are available at http://www.ncbi.nlm.nih.gov/bioproject using bioproject ID: PRJNA1387371. Raw metabolomics mass spectrometry data are available at the MetaboLights (https://www.ebi.ac.uk/metabolights/) data repository under project accession MTBLS13492. Processed MS data is available in **Tables S2-S5**. All other supporting data, including uncropped blots and gels is available in the attached Source Data File.

## Declaration of Interests

The authors declare no competing interests.

## Acknowledgments

This work was supported by a BBSRC award (BB/W014947/1) to LP and CRH, and NIH grant no. AI157247 (A.A.K) and University of Arizona (A.A.K, JTN). This work was also supported by The Royal Society of Edinburgh (SRE-5046) to LP. LF was supported by a Vacation Scholarship from the Microbiology Society. CRH is funded by a Sir Henry Dale Fellowship from the Wellcome Trust and the Royal Society (213455/Z/18/Z). SJF and GGvD are supported by an Australian Research Council Discovery Project (DP230100853). C.B. was supported by the ARISE program that received funding from the European Union’s Horizon 2020 research and innovation programme under the Marie Sklodowska-Curie grant agreement number 945405. The work is supported by the EU project IMAGINE, grant agreement ID: 101094250. We thank the Instrumentation Team of EMBL Grenoble (Gergely Papp & Léa Lecomte) for access to the EasyGrid technology and the European Synchrotron Radiation Facility (ESRF) for provision of synchrotron radiation facilities under proposal ID LS3370 on beamline ID16A.

## Notes

### Competing Interest Statement

The authors have declared no competing interest.

### Summary of Updates

Figure 1 and Figure 5 revised with new data.

## References

1. Chen, L., Min, J. & Wang, F. Copper homeostasis and cuproptosis in health and disease. Sig Transduct Target Ther 7, 378 (2022).

2. Tsvetkov, P. et al. Copper induces cell death by targeting lipoylated TCA cycle proteins. Science 375, 1254–1261 (2022).

3. Besold, A. N., Culbertson, E. M. & Culotta, V. C. The Yin and Yang of Copper During Infection. J Biol Inorg Chem 21, 137–144 (2016).

4. Tsang, T., Davis, C. I. & Brady, D. C. Copper biology. Current Biology 31, R421–R427 (2021).

5. Focarelli, F., Giachino, A. & Waldron, K. J. Copper microenvironments in the human body define patterns of copper adaptation in pathogenic bacteria. PLOS Pathogens 18, e1010617 (2022).

6. Sheldon, J. R. & Skaar, E. P. Metals as phagocyte antimicrobial effectors. Curr Opin Immunol 60, 1–9 (2019).

7. Bollani, L. et al. Congenital Toxoplasmosis: The State of the Art. Front. Pediatr. 10, (2022).

8. Derouin, F. & Pelloux, H. Prevention of toxoplasmosis in transplant patients. Clinical Microbiology and Infection 14, 1089–1101 (2008).

9. Aghabi, D. et al. ZFT is the major iron and zinc transporter in Toxoplasma gondii. eLife 14, RP108666 (2026).

10. Blume, M. et al. Host-derived glucose and its transporter in the obligate intracellular pathogen Toxoplasma gondii are dispensable by glutaminolysis. Proceedings of the National Academy of Sciences 106, 12998–13003 (2009).

11. Dass, S. et al. Toxoplasma LIPIN is essential in channeling host lipid fluxes through membrane biogenesis and lipid storage. Nat Commun 12, 2813 (2021).

12. Parker, K. E. R. et al. The tyrosine transporter of Toxoplasma gondii is a member of the newly defined apicomplexan amino acid transporter (ApiAT) family. PLOS Pathogens 15, e1007577 (2019).

13. Giuliano, C. J. et al. CRISPR-based functional profiling of the Toxoplasma gondii genome during acute murine infection. Nat Microbiol (2024) doi:10.1038/s41564-024-01754-2.

14. Krishnan, A., Kloehn, J., Lunghi, M. & Soldati-Favre, D. Vitamin and cofactor acquisition in apicomplexans: Synthesis *versus* salvage. J Biol Chem 295, 701–714 (2020).

15. Ortiz, J. O., Potter, A. K. & Benmerzouga, I. Novel Drug Targets for the Bradyzoite Form of Toxoplasma gondii. Res Rep Trop Med 16, 25–30 (2025).

16. Maya-Maldonado, K. et al. Mosquito metallomics reveal copper and iron as critical factors for Plasmodium infection. PLOS Neglected Tropical Diseases 15, e0009509 (2021).

17. Rasoloson, D., Shi, L., Chong, C. R., Kafsack, B. F. & Sullivan, D. J. Copper pathways in Plasmodium falciparum infected erythrocytes indicate an efflux role for the copper P-ATPase. Biochem J 381, 803–811 (2004).

18. Pfanner, N., Warscheid, B. & Wiedemann, N. Mitochondrial proteins: from biogenesis to functional networks. Nat Rev Mol Cell Biol 20, 267–284 (2019).

19. MacLean, A. E. et al. Structure, assembly and inhibition of the Toxoplasma gondii respiratory chain supercomplex. Nat Struct Mol Biol 32, 1424–1433 (2025).

20. Maclean, A. E. et al. Complexome profile of Toxoplasma gondii mitochondria identifies divergent subunits of respiratory chain complexes including new subunits of cytochrome bc1 complex. PLOS Pathogens 17, e1009301 (2021).

21. Seidi, A. et al. Elucidating the mitochondrial proteome of Toxoplasma gondii reveals the presence of a divergent cytochrome c oxidase. eLife 7, e38131 (2018).

22. Salman, A. A. & Goldring, J. P. D. Expression and copper binding studies of a Plasmodium falciparum protein with Cox19 copper binding motifs. Exp Parasitol 251, 108572 (2023).

23. Salman, A. A. & Goldring, J. P. D. Expression and copper binding characteristics of Plasmodium falciparum cytochrome c oxidase assembly factor 11, Cox11. Malaria Journal 21, 173 (2022).

24. Huet, D., Rajendran, E., van Dooren, G. G. & Lourido, S. Identification of cryptic subunits from an apicomplexan ATP synthase. Elife 7, e38097 (2018).

25. Leonard, R. A., Tian, Y., Tan, F., Dooren, G. G. van & Hayward, J. A. An essential role for an Fe-S cluster protein in the cytochrome c oxidase complex of Toxoplasma parasites. PLOS Pathogens 19, e1011430 (2023).

26. Painter, H. J., Morrisey, J. M., Mather, M. W. & Vaidya, A. B. Specific role of mitochondrial electron transport in blood-stage Plasmodium falciparum. Nature 446, 88–91 (2007).

27. Harb, O. S. & Roos, D. S. ToxoDB: Functional Genomics Resource for Toxoplasma and Related Organisms. in Toxoplasma gondii: Methods and Protocols (ed. Tonkin, C. J.) 27–47 (Springer US, New York, NY, 2020). doi:10.1007/978-1-4939-9857-9_2.

28. Odberg-Ferragut, C. et al. Molecular cloning, expression analysis and iron metal cofactor characterisation of a superoxide dismutase from Toxoplasma gondii. Mol Biochem Parasitol 106, 121–129 (2000).

29. Sidik, S. M. et al. A Genome-wide CRISPR Screen in Toxoplasma Identifies Essential Apicomplexan Genes. Cell 166, 1423–1435.e12 (2016).

30. Ren, F. et al. X-ray structures of the high-affinity copper transporter Ctr1. Nat Commun 10, 1386 (2019).

31. Wen, M.-H. et al. Elevated intracellular copper induces CTR1 monomerization and prevents copper uptake. Nat Commun 16, 11500 (2025).

32. Clifford, R. J., Maryon, E. B. & Kaplan, J. H. Dynamic internalization and recycling of a metal ion transporter: Cu homeostasis and CTR1, the human Cu+ uptake system. Journal of Cell Science 129, 1711–1721 (2016).

33. Logeman, B. L., Wood, L. K., Lee, J. & Thiele, D. J. Gene duplication and neo-functionalization in the evolutionary and functional divergence of the metazoan copper transporters Ctr1 and Ctr2. J Biol Chem 292, 11531–11546 (2017).

34. Choveaux, D. L., Przyborski, J. M. & Goldring, J. P. D. A Plasmodium falciparum copper-binding membrane protein with copper transport motifs. Malar J 11, 397 (2012).

35. Jiang, J., Nadas, I. A., Kim, M. A. & Franz, K. J. A Mets Motif Peptide Found in Copper Transport Proteins Selectively Binds Cu(I) with Methionine-Only Coordination. Inorg. Chem. 44, 9787–9794 (2005).

36. Kar, S. et al. Copper(II) import and reduction are dependent on His-Met clusters in the extracellular amino terminus of human copper transporter-1. J Biol Chem 298, 101631 (2022).

37. Aller, S. G., Eng, E. T., De Feo, C. J. & Unger, V. M. Eukaryotic CTR Copper Uptake Transporters Require Two Faces of the Third Transmembrane Domain for Helix Packing, Oligomerization, and Function. Journal of Biological Chemistry 279, 53435–53441 (2004).

38. Aghabi, D. et al. The vacuolar iron transporter mediates iron detoxification in Toxoplasma gondii. Nat Commun 14, 3659 (2023).

39. Fairweather, S. J. et al. Coordinated action of multiple transporters in the acquisition of essential cationic amino acids by the intracellular parasite Toxoplasma gondii. PLoS Pathog 17, e1009835 (2021).

40. Shingles, R., Wimmers, L. E. & McCarty, R. E. Copper Transport Across Pea Thylakoid Membranes. Plant Physiol 135, 145–151 (2004).

41. Pfluger, S. L. et al. Receptor for Retrograde Transport in the Apicomplexan Parasite Toxoplasma gondii. Eukaryot Cell 4, 432–442 (2005).

42. Sahu, T. et al. ZIPCO, a putative metal ion transporter, is crucial for Plasmodium liver-stage development. EMBO Molecular Medicine 6, 1387–1397 (2014).

43. Vert, G. et al. IRT1, an Arabidopsis transporter essential for iron uptake from the soil and for plant growth. Plant Cell 14, 1223–1233 (2002).

44. Chasen, N. M., Stasic, A. J., Asady, B., Coppens, I. & Moreno, S. N. J. The Vacuolar Zinc Transporter TgZnT Protects Toxoplasma gondii from Zinc Toxicity. mSphere 4, (2019).

45. Stasic, A. J., Moreno, S. N. J., Carruthers, V. B. & Dou, Z. The Toxoplasma Plant-Like Vacuolar Compartment (PLVAC). J Eukaryot Microbiol 69, e12951 (2022).

46. Pareek, V. & Benkovic, S. Metabolic profiling reveals channeled de novo pyrimidine and purine biosynthesis fueled by mitochondrially generated aspartic acid in cancer cells. Nat Commun 16, 8952 (2025).

47. Fox, B. A. & Bzik, D. J. De novo pyrimidine biosynthesis is required for virulence of Toxoplasma gondii. Nature 415, 926–929 (2002).

48. Bajzikova, M. et al. Reactivation of Dihydroorotate Dehydrogenase-Driven Pyrimidine Biosynthesis Restores Tumor Growth of Respiration-Deficient Cancer Cells. Cell Metabolism 29, 399–416.e10 (2019).

49. Curtabbi, A. et al. Ectopic expression of cytosolic DHODH uncouples de novo pyrimidine biosynthesis from mitochondrial electron transport. Nat Metab 8, 454–466 (2026).

50. Löffler, M., Jöckel, J. & Schuster, G. Dihydroorotat-ubiquinone oxidoreductase links mitochondria in the biosynthesis of pyrimidine nucleotides. Mol Cell Biochem 174, 125–129 (1997).

51. Birth, D., Kao, W.-C. & Hunte, C. Structural analysis of atovaquone-inhibited cytochrome bc1 complex reveals the molecular basis of antimalarial drug action. Nat Commun 5, 4029 (2014).

52. Birsoy, K. et al. An Essential Role of the Mitochondrial Electron Transport Chain in Cell Proliferation Is to Enable Aspartate Synthesis. Cell 162, 540–551 (2015).

53. Sullivan, L. B. et al. Supporting Aspartate Biosynthesis Is an Essential Function of Respiration in Proliferating Cells. Cell 162, 552–563 (2015).

54. Hanse, E. A. et al. Cytosolic malate dehydrogenase activity helps support glycolysis in actively proliferating cells and cancer. Oncogene 36, 3915–3924 (2017).

55. King, E. F. B., Cobbold, S. A., Uboldi, A. D., Tonkin, C. J. & McConville, M. J. Metabolomic Analysis of Toxoplasma gondii Tachyzoites. Methods Mol Biol 2071, 435–452 (2020).

56. MacRae, J. I. et al. Mitochondrial metabolism of glucose and glutamine is required for intracellular growth of Toxoplasma gondii. Cell Host Microbe 12, 682–692 (2012).

57. Hayward, J. A. et al. Real-Time Analysis of Mitochondrial Electron Transport Chain Function in Toxoplasma gondii Parasites Using a Seahorse XFe96 Extracellular Flux Analyzer. Bio Protoc 12, e4288 (2022).

58. Hanna, J. C., Shikha, S., Sloan, M. A. & Harding, C. R. Global translational and metabolic remodeling during iron deprivation in Toxoplasma gondii. mBio 0, e03788–25 (2026).

59. van Dooren, G. G., Yeoh, L. M., Striepen, B. & McFadden, G. I. The Import of Proteins into the Mitochondrion of Toxoplasma gondii. J. Biol. Chem. 291, 19335–19350 (2016).

60. Lacombe, A. et al. Identification of the Toxoplasma gondii mitochondrial ribosome, and characterisation of a protein essential for mitochondrial translation. Mol Microbiol 112, 1235–1252 (2019).

61. Waldman, B. S. et al. Identification of a Master Regulator of Differentiation in Toxoplasma. Cell 180, 359–372.e16 (2020).

62. Merritt, E. F. et al. Transcriptional Profiling Suggests T Cells Cluster around Neurons Injected with Toxoplasma gondii Proteins. mSphere 5, e00538–20 (2020).

63. Tomita, T. et al. The Toxoplasma gondii cyst wall protein CST1 is critical for cyst wall integrity and promotes bradyzoite persistence. PLoS Pathog 9, e1003823 (2013).

64. Kochanowsky, J. A., Chandrasekaran, S., Sanchez, J. R., Thomas, K. K. & Koshy, A. A. ROP16-mediated activation of STAT6 enhances cyst development of type III Toxoplasma gondii in neurons. PLoS Pathog 19, e1011347 (2023).

65. Cabral, C. M. et al. Neurons are the Primary Target Cell for the Brain-Tropic Intracellular Parasite Toxoplasma gondii. PLoS Pathog 12, e1005447 (2016).

66. Kropat, J. et al. A regulator of nutritional copper signaling in Chlamydomonas is an SBP domain protein that recognizes the GTAC core of copper response element. Proc Natl Acad Sci U S A 102, 18730–18735 (2005).

67. Liu, L., Qi, J., Yang, Z., Peng, L. & Li, C. Low-affinity copper transporter *CTR2* is regulated by copper-sensing transcription factor Mac1p in *Saccharomyces cerevisiae*. Biochemical and Biophysical Research Communications 420, 600–604 (2012).

68. Rutherford, J. C. & Bird, A. J. Metal-Responsive Transcription Factors That Regulate Iron, Zinc, and Copper Homeostasis in Eukaryotic Cells. Eukaryot Cell 3, 1–13 (2004).

69. Zhang, D.-L., Hughes, R. M., Ollivierre-Wilson, H., Ghosh, M. C. & Rouault, T. A. A ferroportin transcript that lacks an iron-responsive element enables duodenal and erythroid precursor cells to evade translational repression. Cell Metab 9, 461–473 (2009).

70. Kato, J. et al. Iron/IRP-1-dependent regulation of mRNA expression for transferrin receptor, DMT1 and ferritin during human erythroid differentiation. Experimental Hematology 35, 879–887 (2007).

71. Sloan, M. A., Scott, A., Aghabi, D., Mrvova, L. & Harding, C. R. Iron-mediated post-transcriptional regulation in Toxoplasma gondii. PLOS Pathogens 21, e1012857 (2025).

72. Pena, M. M., Puig, S. & Thiele, D. J. Characterization of the Saccharomyces cerevisiae high affinity copper transporter Ctr3. J Biol Chem 275, 33244–33251 (2000).

73. Cai, Z. et al. Cu-sensing transcription factor Mac1 coordinates with the Ctr transporter family to regulate Cu acquisition and virulence in Aspergillus fumigatus. Fungal Genet Biol 107, 31–43 (2017).

74. Collins, J. F., Prohaska, J. R. & Knutson, M. D. Metabolic crossroads of iron and copper. Nutr Rev 68, 133–147 (2010).

75. Strenkert, D. et al. Distinct function of Chlamydomonas CTRA-CTR transporters in Cu assimilation and intracellular mobilization. Metallomics. 16, mfae013 (2024).

76. Barylyuk, K. et al. A Comprehensive Subcellular Atlas of the Toxoplasma Proteome via hyperLOPIT Provides Spatial Context for Protein Functions. Cell Host & Microbe 28, 752–766.e9 (2020).

77. Curnock, R. & Cullen, P. J. Mammalian copper homeostasis requires retromer-dependent recycling of the high-affinity copper transporter 1. J Cell Sci 133, jcs249201 (2020).

78. Krishnan, A. et al. Functional and Computational Genomics Reveal Unprecedented Flexibility in Stage-Specific Toxoplasma Metabolism. Cell Host & Microbe 27, 290–306.e11 (2020).

79. Buchet, K. & Godinot, C. Functional F1-ATPase essential in maintaining growth and membrane potential of human mitochondrial DNA-depleted rho degrees cells. J Biol Chem 273, 22983–22989 (1998).

80. Kunzová, M., Doleželová, E., Moos, M., Panicucci, B. & Zíková, A. Reversal of ATP synthase is a key attribute accompanying cellular differentiation of Trypanosoma brucei insect forms. Commun Biol 9, 680 (2026).

81. Nývltová, E., Dietz, J. V., Seravalli, J., Khalimonchuk, O. & Barrientos, A. Coordination of metal center biogenesis in human cytochrome c oxidase. Nat Commun 13, 3615 (2022).

82. Keyhani, E. & Keyhani, J. Cytochrome *c* oxidase biosynthesis and assembly in *Candida utilis* yeast cells. Archives of Biochemistry and Biophysics 167, 596–602 (1975).

83. Secic, D. et al. Dynamic assessment of the allocation of copper to cytochrome c oxidase using size-exclusion chromatography (SEC) combined with inductively coupled plasma mass spectrometry (ICP-MS). Journal of Biological Chemistry 111278 (2026) doi:10.1016/j.jbc.2026.111278.

84. Bertinato, J., Swist, E., Plouffe, L. J., Brooks, S. P. J. & L’abbé, M. R. Ctr2 is partially localized to the plasma membrane and stimulates copper uptake in COS-7 cells. Biochem J 409, 731–740 (2008).

85. Rees, E. M., Lee, J. & Thiele, D. J. Mobilization of Intracellular Copper Stores by the Ctr2 Vacuolar Copper Transporter. Journal of Biological Chemistry 279, 54221–54229 (2004).

86. Buchholz, K. R. et al. Identification of tissue cyst wall components by transcriptome analysis of in vivo and in vitro Toxoplasma gondii bradyzoites. Eukaryot Cell 10, 1637–1647 (2011).

87. Fux, B. et al. Toxoplasma gondii strains defective in oral transmission are also defective in developmental stage differentiation. Infect Immun 75, 2580–2590 (2007).

88. Hennigar, S. R. & McClung, J. P. Nutritional Immunity: Starving Pathogens of Trace Minerals. Am J Lifestyle Med 10, 170–173 (2016).

89. Monteith, A. J. & Skaar, E. P. The impact of metal availability on immune function during infection. Trends Endocrinol Metab 32, 916–928 (2021).

90. Christiansen, C. et al. In vitro maturation of Toxoplasma gondii bradyzoites in human myotubes and their metabolomic characterization. Nat Commun 13, 1168 (2022).

91. Shukla, A. et al. Glycolysis is important for optimal asexual growth and formation of mature tissue cysts by Toxoplasma gondii. Int J Parasitol 48, 955–968 (2018).

92. Amos, B. et al. VEuPathDB: the eukaryotic pathogen, vector and host bioinformatics resource center. Nucleic Acids Research 50, D898–D911 (2022).

93. Madeira, F. et al. The EMBL-EBI Job Dispatcher sequence analysis tools framework in 2024. Nucleic Acids Res 52, W521–W525 (2024).

94. Letunic, I. & Bork, P. Interactive Tree Of Life (iTOL) v5: an online tool for phylogenetic tree display and annotation. Nucleic Acids Res 49, W293–W296 (2021).

95. Pfaffl, M. W. A new mathematical model for relative quantification in real-time RT–PCR. Nucleic Acids Res 29, e45 (2001).

96. Rajendran, E. et al. Cationic amino acid transporters play key roles in the survival and transmission of apicomplexan parasites. Nat Commun 8, 14455 (2017).

97. Wagner, C. A., Friedrich, B., Setiawan, I., Lang, F. & Bröer, S. The use of Xenopus laevis oocytes for the functional characterization of heterologously expressed membrane proteins. Cell Physiol Biochem 10, 1–12 (2000).

98. Bröer, S. Xenopus laevis Oocytes. Methods Mol Biol 637, 295–310 (2010).

99. Cinquetti, R. et al. Characterization of Transport Activity of SLC11 Transporters in Xenopus laevis Oocytes by Fluorophore Quenching. SLAS Discov 26, 798–810 (2021).

100. Shen, B., Brown, K. M., Lee, T. D. & Sibley, L. D. Efficient Gene Disruption in Diverse Strains of Toxoplasma gondii Using CRISPR/CAS9. mBio 5, 10.1128/mbio.01114-14 (2014).

101. Caffaro, C. E. et al. A Nucleotide Sugar Transporter Involved in Glycosylation of the Toxoplasma Tissue Cyst Wall Is Required for Efficient Persistence of Bradyzoites. PLoS Pathog 9, e1003331 (2013).

102. Messina, M., Niesman, I., Mercier, C. & Sibley, L. D. Stable DNA transformation of *Toxoplasma gondii* using phleomycin selection. Gene 165, 213–217 (1995).

103. Curt-Varesano, A., Braun, L., Ranquet, C., Hakimi, M.-A. & Bougdour, A. The aspartyl protease TgASP5 mediates the export of the Toxoplasma GRA16 and GRA24 effectors into host cells. Cell Microbiol 18, 151–167 (2016).

104. Soldati, D. & Boothroyd, J. C. Transient Transfection and Expression in the Obligate Intracellular Parasite Toxoplasma gondii. Science 260, 349–352 (1993).

105. van de Ven, E., Melchers, W., Galama, J., Camps, W. & Meuwissen, J. Identification of Toxoplasma gondii infections by BI gene amplification. J Clin Microbiol 29, 2120–2124 (1991).

106. Saeij, J. P. J., Arrizabalaga, G. & Boothroyd, J. C. A Cluster of Four Surface Antigen Genes Specifically Expressed in Bradyzoites, SAG2CDXY, Plays an Important Role in Toxoplasma gondii Persistence. Infect Immun 76, 2402–2410 (2008).

107. Kim, S.-K., Karasov, A. & Boothroyd, J. C. Bradyzoite-specific surface antigen SRS9 plays a role in maintaining Toxoplasma gondii persistence in the brain and in host control of parasite replication in the intestine. Infect Immun 75, 1626–1634 (2007).

108. Jacobs, K., Charvat, R. & Arrizabalaga, G. Identification of Fis1 Interactors in Toxoplasma gondii Reveals a Novel Protein Required for Peripheral Distribution of the Mitochondrion. mBio 11, e02732–19 (2020).

109. Breinich, M. S. et al. A dynamin is required for the biogenesis of secretory organelles in Toxoplasma gondii. Curr Biol 19, 277–286 (2009).

110. Nagamune, K., Beatty, W. L. & Sibley, L. D. Artemisinin induces calcium-dependent protein secretion in the protozoan parasite Toxoplasma gondii. Eukaryot Cell 6, 2147–2156 (2007).

111. Bissardon, C. et al. Cell Culture on Silicon Nitride Membranes and Cryopreparation for Synchrotron X-ray Fluorescence Nano-analysis. J Vis Exp 10.3791/60461 (2019) doi:10.3791/60461.

112. Gemin, O. et al. EasyGrid: a versatile platform for automated cryo-EM sample preparation and quality control. 2024.01.18.576170 Preprint at 10.1101/2024.01.18.576170 (2024).

113. Silva, J. C. da et al. Efficient concentration of high-energy x-rays for diffraction-limited imaging resolution. Optica, OPTICA 4, 492–495 (2017).

114. Villar, F. et al. Nanopositioning for the ESRF ID16A Nano-Imaging Beamline. Synchrotron Radiation News 31, 9–14 (2018).

115. Utica, G., et al. ARDESIA-16: a 16-channel SDD-based spectrometer for energy dispersive X-ray fluorescence spectroscopy. J. Inst. 16, P07057 (2021).

116. Solé, V. A., Papillon, E., Cotte, M., Walter, Ph. & Susini, J. A multiplatform code for the analysis of energy-dispersive X-ray fluorescence spectra. Spectrochimica Acta Part B: Atomic Spectroscopy 62, 63–68 (2007).

117. Bolger, A. M., Lohse, M. & Usadel, B. Trimmomatic: a flexible trimmer for Illumina sequence data. Bioinformatics 30, 2114–2120 (2014).

118. Kim, D., Langmead, B. & Salzberg, S. L. HISAT: a fast spliced aligner with low memory requirements. Nat Methods 12, 357–360 (2015).

119. Love, M. I., Huber, W. & Anders, S. Moderated estimation of fold change and dispersion for RNA-seq data with DESeq2. Genome Biology 15, 550 (2014).

120. Degasperi, A. et al. Evaluating Strategies to Normalise Biological Replicates of Western Blot Data. PLOS ONE 9, e87293 (2014).

121. Pountain, A. W. & Barrett, M. P. Untargeted metabolomics to understand the basis of phenotypic differences in amphotericin B-resistant Leishmania parasites. Wellcome Open Res 4, 176 (2019).

122. Creek, D. J., Jankevics, A., Burgess, K. E. V., Breitling, R. & Barrett, M. P. IDEOM: an Excel interface for analysis of LC-MS-based metabolomics data. Bioinformatics 28, 1048–1049 (2012).

123. Scheltema, R. A., Jankevics, A., Jansen, R. C., Swertz, M. A. & Breitling, R. PeakML/mzMatch: A File Format, Java Library, R Library, and Tool-Chain for Mass Spectrometry Data Analysis. Anal. Chem. 83, 2786–2793 (2011).

124. Salek, R. M., Steinbeck, C., Viant, M. R., Goodacre, R. & Dunn, W. B. The role of reporting standards for metabolite annotation and identification in metabolomic studies. Gigascience 2, 13 (2013).

125. Pang, Z. et al. MetaboAnalystR 4.0: a unified LC-MS workflow for global metabolomics. Nat Commun 15, 3675 (2024).

126. Su, X., Lu, W. & Rabinowitz, J. D. Metabolite Spectral Accuracy on Orbitraps. Anal Chem 89, 5940–5948 (2017).

127. Ritchie, M. E. et al. limma powers differential expression analyses for RNA-sequencing and microarray studies. Nucleic Acids Res 43, e47 (2015).

128. Kochanowsky, J. A., Chandrasekaran, S., Sanchez, J. R., Thomas, K. K. & Koshy, A. A. ROP16-mediated activation of STAT6 enhances cyst development of type III Toxoplasma gondii in neurons. PLOS Pathogens 19, e1011347 (2023).

129. Koshy, A. A. et al. Toxoplasma Co-opts Host Cells It Does Not Invade. PLOS Pathogens 8, e1002825 (2012).

